# Target-induced Argonaute-HNH filaments confer bacterial immunity

**DOI:** 10.1101/2025.04.06.647449

**Authors:** Anna Kanevskaya, Lidiya Lisitskaya, Andrey V. Moiseenko, Olga S. Sokolova, Nikolai N. Sluchanko, Andrey Kulbachinskiy

## Abstract

Argonaute proteins provide innate immunity in all domains of life through guide-dependent recognition of invader nucleic acids. While eukaryotic Argonautes (eAgos) act on RNA during RNA interference, prokaryotic Argonautes (pAgos) mainly recognize DNA targets. Many eAgos and some pAgos are active nucleases that directly cleave their targets. In contrast, short pAgos lack the nuclease activity and are co-encoded with additional effectors. Diverse effector domains in short pAgo systems include NADases and tentative nucleases, but their mechanisms of activation remain largely unknown. Here, we characterize SPARHA systems (<u>s</u>hort <u>p</u>rokaryotic argonautes, <u>H</u>NH-<u>a</u>ssociated) encoding HNH nuclease effectors. We show that short pAgo activates the HNH effector after RNA-guided DNA recognition. Target recognition induces formation of filaments of SPARHA with double active sites formed at the interfaces of repetitive HNH domains, which results in indiscriminate collateral degradation of DNA and protects bacterial population from invaders. The results show that pAgos and associated effectors act as modular two-component systems that translate recognition of specific DNA into immune response through assembly of supramolecular complexes, deleterious for invaders and potentially useful for biotechnology.

---

Argonaute proteins were first identified as bona fide nucleases with a ‘slicer’ activity in the core RNA interference machinery involved in gene silencing in eukaryotes. Eukaryotic argonautes (eAgos) are programmed with small guide RNAs of different classes loaded through specialized pathways and recognize viral, transposonic or cellular RNA transcripts leading to their degradation, inhibition of translation, or transcriptional repression ^1-6^. In contrast, most prokaryotic argonautes (pAgos) recognize DNA targets, using either guide DNAs or guide RNAs ^7-15^. This correlates with their proposed role in cell defense against invader genetic elements and phages, most of which have DNA genomes. Small groups of RNA-targeting pAgos were discovered recently, highlighting the diversity of pAgo functions and their potential involvement in additional regulatory pathways ^16-18^. Unlike eAgos that always contain six conserved domains (N, L1, PAZ, L2, MID and PIWI), the majority of pAgos are ‘short’ and contain only two of these domains, MID and PIWI, always with substitutions in PIWI inactivating their nuclease activity ^3,5,7,19,20^. Instead, short pAgos are co-encoded with partner proteins containing an additional effector domain fused to an APAZ domain ^3,19,20^. APAZ corresponds to the N-terminal, L1 and L2 domains of full-length argonautes and forms a complex with short pAgo, thus bringing the effector domain under the control of pAgo ^19,21-24^. Short pAgos systems form four major clades associated with different effectors: SIR2 domains in clades S1A and S1B (encoded separately or fused to pAgos), TIR domains in clade S2A, and various effectors in clade S2B, including tentative nucleases of several classes (DREN, Mrr-like and HNH) and additional uncharacterized effectors (Fig. 1a).

**Figure 1.**
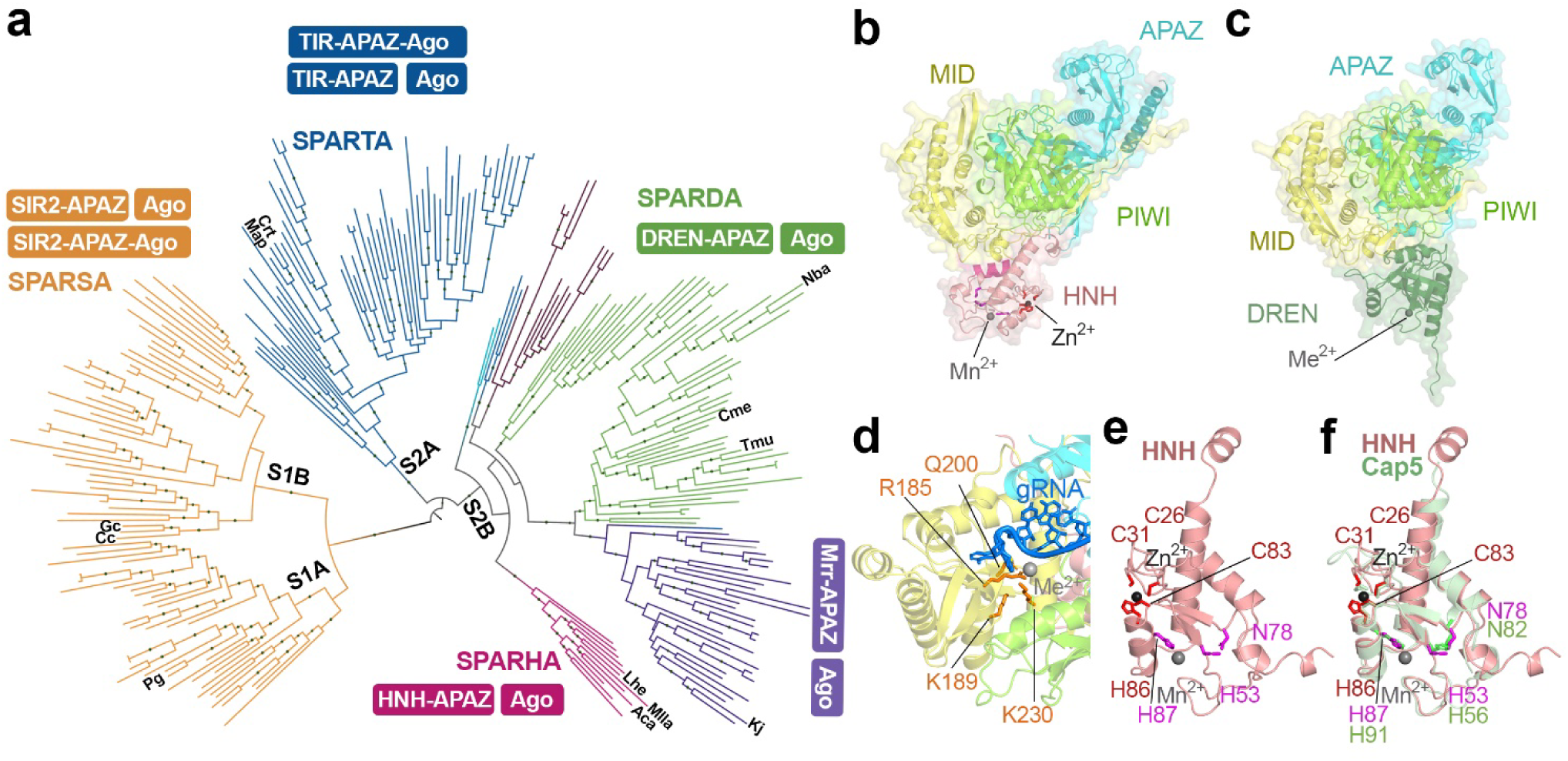
Phylogeny and structure of SPARHA systems. **a**, Phylogenetic tree and operon structure of short pAgo systems. Previously studied short pAgos ^19,21,22,24^ and pAgos analyzed in this study are indicated. The dots represent bootstrap values >75%. **b,** AlphaFold 3 (AF3) model of MllaSPARHA, with an HNH effector nuclease. **c,** AF3 model of CmeSPARDA, with a DREN effector nuclease ^24^. **d,** Guide-binding pocket in the MID domain of MllaAgo. **e,** Modelled structure of the HNH effector of MllaSPARHA. Catalytic HNH residues holding a Mn^2+^ ion and cysteine/histidine residues holding a Zn^2+^ ion are shown in magenta and red, respectively. **f,** Superimposition of HNH domains in MllaSPARHA (AF3 model) and Cap5 (PDB: 8fmf)^27^.

Pioneering studies of short pAgo systems demonstrated that guide-RNA-dependent recognition of target DNA by short pAgos activates associated SIR2 and TIR effectors, causing depletion of NAD^+^/NADH (SPARSA and SPARTA systems). We and others recently showed that short pAgos associated with the DREN domain (SPARDA systems) can indiscriminately degrade DNA and RNA upon target DNA recognition ^23,24^. These activities inhibit plasmid transformation and phage replication by an abortive infection mechanism, leading to cell death or dormancy ^19,22,24,25^. However, the mechanisms of activation of pAgo-associated nucleases have not been studied in detail.

Here, we have investigated a distinct clade of S2B pAgo systems encoding putative HNH nucleases (SPARHA, for <u>s</u>hort <u>p</u>rokaryotic <u>ar</u>gonautes associated with <u>H</u>NH-<u>A</u>PAZ) and demonstrated that the HNH nuclease is activated by its pAgo partner after guide-dependent target DNA recognition. Activated SPARHA forms protein-nucleic acid filaments with robust nuclease activity, degrading cellular DNA and protecting bacterial population from invaders by abortive infection. Analysis of SPARHA provides a striking example of a filamentous immune nuclease and shows that short pAgos can serve as an assembly platform for higher-order protein-nucleic acid complexes in prokaryotic immunity.

## SPARHA is a short pAgo system with an HNH effector

To investigate the activities of SPARHA systems, we selected MllaAgo from the longest-named myxobacterium *Myxococcus llanfairpwllgwyngyllgogerychwyrndrobwllllantysiliogogogochensis*, AcaAgo from a biomining gammaproteobacterium *Acidithiobacillus caldus* and LheAgo from a Muppet-looking cyanobacterium *Leptolyngbya sp.* ‘hensonii’, all belonging to the HNH clade of short pAgos (Fig. 1a). Structural modeling of MllaSPARHA and AcaSPARHA showed that short pAgos form heterodimeric complexes with their associated HNH-APAZ effectors, with the MID, PIWI and APAZ domains forming a characteristic bilobal structure similar to previously characterized long and short pAgos and involved in guide binding and target recognition (Fig. 1b, Fig. ED11; compare with the SPARDA complex in Fig. 1c). Similarly to other short pAgos, the PIWI domain lacks the catalytic tetrad residues, while the MID domain contains a conserved RKQK motif involved in 5’-guide binding (Fig. 1d, Extended Data Fig. ED1a). The HNH domain is located outside of the nucleic acid binding cleft formed by pAgo-APAZ suggesting that it may be involved in collateral nucleic acid cleavage (Fig. 1b). It has a typical fold of His-Me finger nucleases, with two catalytic His residues located on adjacent β strand and α helix (Fig. 1e,f, Fig. ED1b)^26^. The HNH domain of SPARHA can be perfectly superimposed on the HNH domain of a CBASS effector Cap5 (Fig. 1f) ^27^, which is an active DNA nuclease. We cloned and expressed selected SPARHA systems in *Escherichia coli* and purified them to homogeneity. All three pAgos were co-purified with their HNH-APAZ partners at all chromatography steps, confirming that they form a stable complex (Fig. ED1c,d,e).

## SPARHA is activated upon target DNA recognition

To test the nuclease activity of SPARHA, we loaded MllaSPARHA with short guide RNAs or DNAs, supplied it with complementary single-stranded RNA or DNA targets and incubated with various collateral nucleic acid substrates (Fig. 2a). MllaSPARHA degraded short linear dsDNA in the presence of guide RNA and target DNA, but was inactive in the absence of guide RNA or with the other three guide/target combinations (Fig. 2b). The observed specificity of SPARHA for RNA guides and DNA targets corresponds to previously characterized short pAgo systems ^19,22-24^.

**Figure 2.**
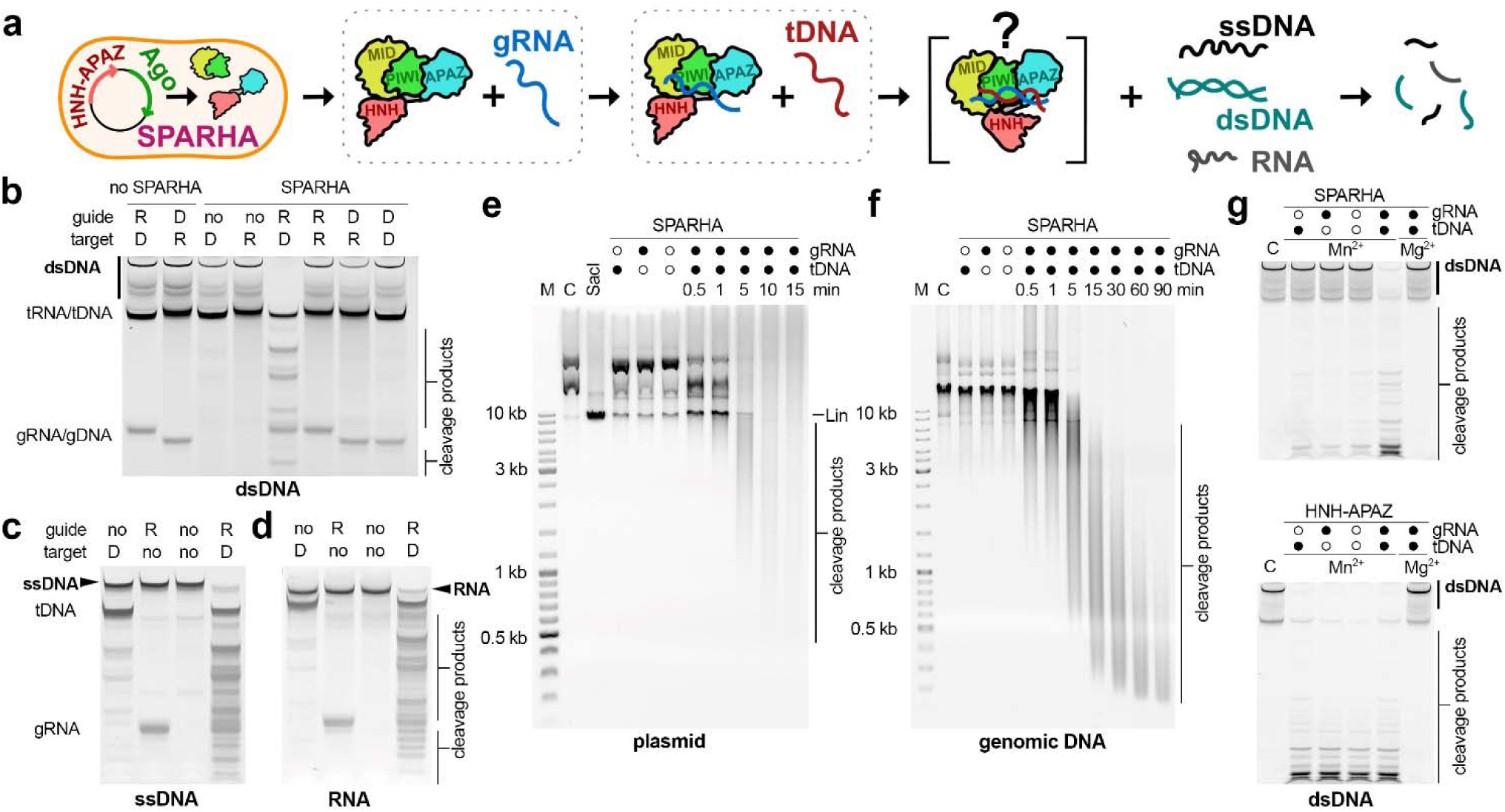
Collateral nuclease activity of SPARHA. **a,** Scheme of the assay. **b-d**, Collateral activity of MllaSPARDA with linear dsDNA (b), ssDNA (c) and RNA (d) substrates. The reactions contained RNA (R) or DNA (D) guides and targets. The dsDNA substrate migrates as several bands due to partial denaturation during PAGE. **e-f,** Kinetics of cleavage of plasmid (e) and genomic (f) DNA by MllaSPARHA. **g,** Cleavage of linear dsDNA by LheSPARHA (top) and LheHNH-APAZ (bottom). Representative gels from three independent experiments, which produced similar results, are shown. Positions of collateral substrates, guides (gRNA/gDNA), targets (tRNA/tDNA) and cleavage products are indicated. Panels b-f, SYBR Gold staining; panel g, fluorescence of HEX-labeled dsDNA substrate. C, control samples without SPARHA; M, length markers; Lin, linear plasmid, treated with SacI. See Fig. ED2 and Table S2 for guide, target and substrate sequences. Panel **a** created with Inkscape (http://www.inkscape.org).

When loaded with guide RNA and target DNA, MllaSPARHA also cleaved ssDNA and RNA oligonucleotide substrates (Fig. 2c,d), but with lower rates in comparison with dsDNA of an identical sequence (Fig. ED2b). MllaSPARHA could digest a ten-fold excess of collateral substrates indicating that it is a multiple-turnover enzyme (Fig. ED2b). MllaSPARHA gradually degraded purified plasmid DNA and genomic DNA into short fragments (Fig. 2e,f), showing that it can introduce multiple cleavages in the same molecule. Similarly, AcaSPARHA and LheSPARHA effectively degraded linear dsDNA and plasmid DNA, when loaded with RNA guides and complementary DNA targets, but were much less active with RNA substrates (Fig. ED3a,b,c,i,j,k). Therefore, SPARHA systems preferentially act on dsDNA targets.

The nuclease activity of SPARHA systems was observed in the presence of Mn^2+^ but not Mg^2+^, Zn^2+^, Co^2+^ or Ni^2+^ ions (Fig. ED2c and Fig. ED3d for MllaSPARHA and AcaSPARHA); such preference was rarely observed in previously studied HNH nucleases ^28^. Substitutions of predicted catalytic residues in the HNH domain in MllaSPARHA (H53A) and AcaSPARHA (N71A/H80A) completely abolished their nuclease activity (Fig. ED2d and Fig. ED3e). Both MllaSPARHA and AcaSPARHA were optimally active at physiological temperatures, 25-45 °C and 30-50 °C, respectively (Fig. ED2e and Fig. ED3f). MllaSPARHA was active at 5-100 mM KCl, while AcaSPARHA became inactive at >50 mM KCl (Fig. ED2f and Fig. ED3g). MllaSPARHA could use guide RNAs of various lengths (16-24 nt) (Fig. ED2g), and was active with guides containing or lacking the 5’-phosphate (Fig. ED2h) or guides containing any 5’-nucleotide (Fig. ED2i). In comparison, AcaSPARHA preferred 5’-A guide RNAs, while LheSPARHA was most active with a 5’-UU guide RNA (Fig. ED3h,i).

To reveal the role of pAgo in the regulation of the HNH nuclease, we compared nuclease activities of free and pAgo-bound HNH effector. While LheSPARHA was only active with guide RNA and target DNA, isolated LheHNH-APAZ efficiently degraded collateral dsDNA independently of the presence of guides and targets (Fig. 2g, Fig. ED3l). Therefore, the HNH effector is kept inactive in the complex with pAgo, while target recognition by pAgo likely induces structural changes of SPARHA, unleashing its nuclease activity.

Collateral DNA cleavage by SPARHA can potentially be used for DNA detection in various applications. As a proof of principle, we designed a fluorometric assay in which recognition of specific DNA by SPARHA results in cleavage of a dsDNA beacon containing a fluorophore and a quencher, thereby generating a fluorescence signal (Fig. ED4a). The concentration limit of DNA detection in this assay using AcaSPARHA was ∼ 3 nM (Fig. ED4b). The sensitivity of SPARHA was lower than that of SPARDA ^24^, due to higher background activity in the absence of target DNA (Fig. ED4b,c), but higher than that of SPARTA, which generates fluorescent signal by cleaving NAD^+^ derivatives (Fig. ED4d) ^19^.

## SPARHA is activated by plasmids and phages

We next tested whether SPARHA can provide protection against invaders, plasmids or phages. We expressed MllaSPARHA or its mutant variant with substitutions in the HNH nuclease (CD, catalytically dead) from a pBAD vector in *E. coli* either lacking or containing an additional ‘interfering’ plasmid (pCDF or pACYC), measured growth rates in liquid cultures and determined the numbers of live cells (CFU, colony forming units) in these cultures (Fig. 3a). Expression of wild-type but not CD MllaSPARHA induced lysis of cell culture at ∼7.5 hours of growth in the presence of pCDF (Fig. 3b, right). Arrest of cell growth by wild-type MllaSPARHA was also observed with the pACYC plasmid (Fig. ED5a). No changes in cell growth in these conditions were observed in the absence of the interfering plasmid (Fig. 3b, left). AcaSPARHA also delayed cell growth in the presence but not in the absence of pCDF, although its effects were weaker than those of MllaSPARHA (Fig. ED5b). LheSPARHA almost completely prevented cell growth in the presence of pCDF, but was toxic even in its absence (Fig. ED5b). We therefore focused on further analysis of MllaSPARHA. In the presence of pCDF, CFU numbers measured just before the onset of cell lysis (∼8 h post induction) were dramatically decreased in cultures expressing wild-type but not CD MllaSPARHA, relative to an empty pBAD control (Fig. 3c, Fig. ED5c). Similar, albeit weaker, effects on CFU numbers were observed in cell cultures containing the pACYC plasmid (Fig. ED5d). No changes in CFU numbers were observed in the absence of interfering plasmids. Similar effects of the growth rate and CFU numbers were observed at different temperatures, although pCDF-induced cell lysis was delayed at a higher temperature (Fig. ED5e). Identical drops in the CFU numbers were observed with antibiotics corresponding to the expression vector (pBAD, Amp^R^) or the invader plasmid (pCDF, Sm^R^ or pACYC, Cm^R^), or when the cells were grown in the absence of antibiotics (Fig. 3с, Fig. ED5d,e,f). This indicated that the observed CFU changes resulted from an overall decrease in live cell numbers and not from selective elimination of any of the plasmids from the culture.

**Figure 3.**
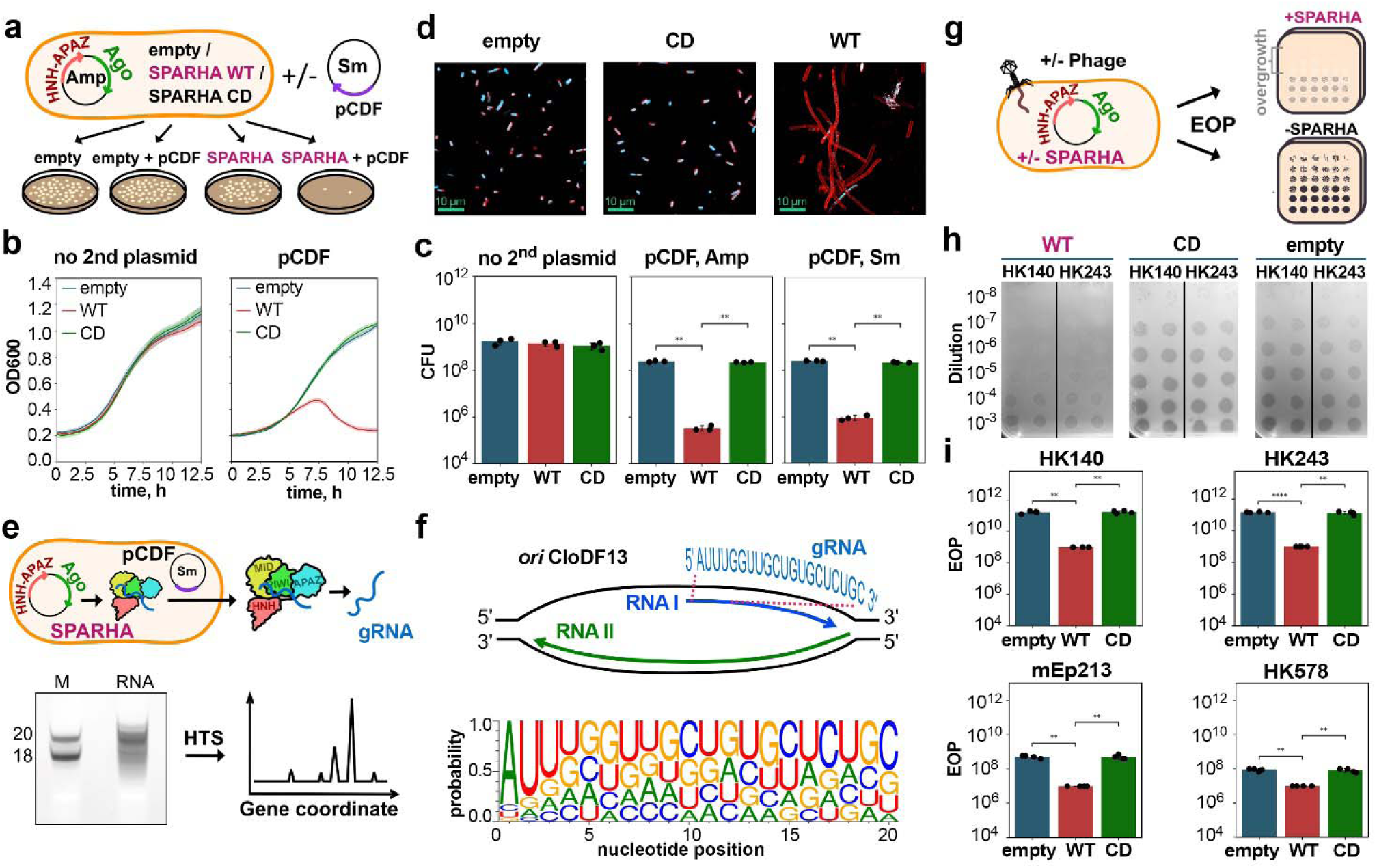
The antiplasmid and antiphage activity of SPARHA. **a,** Plasmid interference assay. **b,** Growth of *E. coli* cells expressing wild-type (WT) or catalytically-dead (CD) MllaSPARHA or containing empty pBAD, in the absence and in the presence of the interfering plasmid (pCDF). **c,** CFU numbers in cell cultures expressing WT or CD MllaSPARHA or containing empty pBAD, depending on the presence of pCDF, collected at 8 h of growth. CFU were measured on LB plates containing Amp (for experiments with and without pCDF) or Sm (for experiments with pCDF). Means and standard deviations from 3 independent experiments. Statistically significant changes are (from left to right: **P=0.0035, **P=0.0013, **P=0.001, **P=0.0014**, calculated using the two-sided t-test for independent samples with Bonferroni correction). (d) Staining of *E. coli* cells expressing WT or CD MllaSPARHA, or containing empty pBAD, in the presence of the interfering plasmid (pCDF) with DNA-specific (cyan) and membrane-specific (red) dyes. Cell cultures were collected at 7.5 hours of growth. The scale bar is 10 μm. Representative fields of view from 3 independent experiments (see also Fig. ED5h). **e,** Scheme of small RNA purification (top), their analysis by PAGE and sequencing (bottom). **f,** Scheme of the CloDF13 *ori* region of pCDF(top) and nucleotide logo of small guide RNAs bound to MllaSPARHA in *E. coli* (bottom). **g,** Phage infection assay. **h,i** Analysis of the effects of WT or CD MllaSPARHA, in comparison with empty pBAD, on the EOP for selected phages. Serial dilutions of phage stocks are indicated on the left (h). The EOP values are averages and standard deviations from 4 independent experiments (i). Statistically significant changes are indicated (from left to right, top to bottom: **P=0.0014, **P=0.0017, ***P=0.00008, **P=0.0033, **P=0.0017, **P=0.0062, **P=0.0015, **P=0.0059**; calculated using the two-sided t-test for independent samples with Holm-Bonferroni correction). Panels **a, e** and **g** were created with Inkscape (http://www.inkscape.org).

To visualize these changes, cells expressing MllaSPARHA and containing an invader plasmid (pCDF) were collected just before the onset of culture lysis and stained with fluorescent dyes specific for cell membranes and DNA. While the control culture and the culture with CD MllaSPARHA mostly contained cells with normal morphology and DNA content, expression of wild-type MllaSPARHA induced formation of long cell filaments lacking DNA (Fig. 3d and Fig. ED5h). Therefore, the activation of MllaSPARHA in the presence of pCDF results in extensive degradation of cellular DNA and causes cell filamentation, possibly by inducing SOS response preventing cell division.

To understand the mechanism of SPARHA activation by the invader plasmid, we isolated and sequenced small guide RNAs bound to MllaSPARHA in cell cultures collected before lysis (Fig. 3e, Fig. ED6a). Most guide RNAs were 17-21 nt long (Fig. ED6b) and had a conserved AU dinucleotide at their 5’-ends (Fig. 3f). Guide RNAs were highly enriched with reads corresponding to plasmid transcripts (34.8% to pCDF, 22.6% to pBAD, 42.6% to chromosomal DNA, average from two replicates; Table S1). The most overrepresented guide RNA exactly corresponded to the 5’-part of a regulatory RNA I from the CloDF13 origin of replication of pCDF (12.2% of ≥20 nt reads in the library) (Fig. 3f, Fig. ED6c). Other parts of the pCDF and pBAD plasmids also generated abundant RNAs bound by MllaSPARHA (Fig. ED6c-e). Similar preferences for guide RNAs from the plasmid *ori* were observed for AcaSPARHA (Fig. ED6b,c). Therefore, SPARHA systems are likely activated by guide RNAs produced from *ori*-specific plasmid transcripts.

Finally, we tested the effects of MllaSPARHA on phage infection (Fig. 3g). Expression of wild-type MllaSPARHA decreased the efficiency of plating (EOP) for 4 out of 13 tested phages, HK140, HK243, mEp213 and HK578 (by 2-3 orders of magnitude), in comparison with CD MllaSPARHA or an empty pBAD control (Fig. 3h,i and Fig. ED7a). No effects were observed for P1, λ, T5, T7 and other phages (Fig. ED7b). Similar effects on sensitive phages were observed at 25°C and 30°C, although changes in EOP were less pronounced at the higher temperature (compare Fig. 3h and Fig. ED7a with Fig. ED7c). Intriguingly, phage spots were initially formed even with sensitive phages (Fig. ED7d) but were later overgrown with cell lawn (Fig. 3g,h), suggesting that SPARHA might be optimally activated at later stages of cell growth.

## SPARHA is activated by filament formation

To understand the mechanism of DNA processing by the activated SPARHA complex, we mapped the sites of cleavage by MllaSPARHA in the chromosomal and plasmid DNA of *E. coli*. The DNA samples were treated with activated MllaSPARHA, producing ∼300-4000 bp fragments, the cleavage sites were blunted, retaining intact 5’-ends, ligated with adaptors and sequenced (Fig. 4a, Fig. ED8a). Mapping of the cleaved 5’-ends on the reference genome showed that they were frequently located at a one-nucleotide distance opposite each other (Fig. 4b), corresponding to double-stranded breaks (DSBs) with 1 nt 3’-overhangs. Sequence logo revealed a strong preference for a (G/C/A)T dinucleotide at both 5’-ends of the DSB, in both genomic and plasmid DNA (Fig. 4c, Fig. ED8b). No such pattern was observed in control samples not treated with MllaSPARHA (Fig. ED8b). This indicates that DNA cleavage is performed by a dimeric nuclease site, with limited sequence specificity.

**Figure 4.**
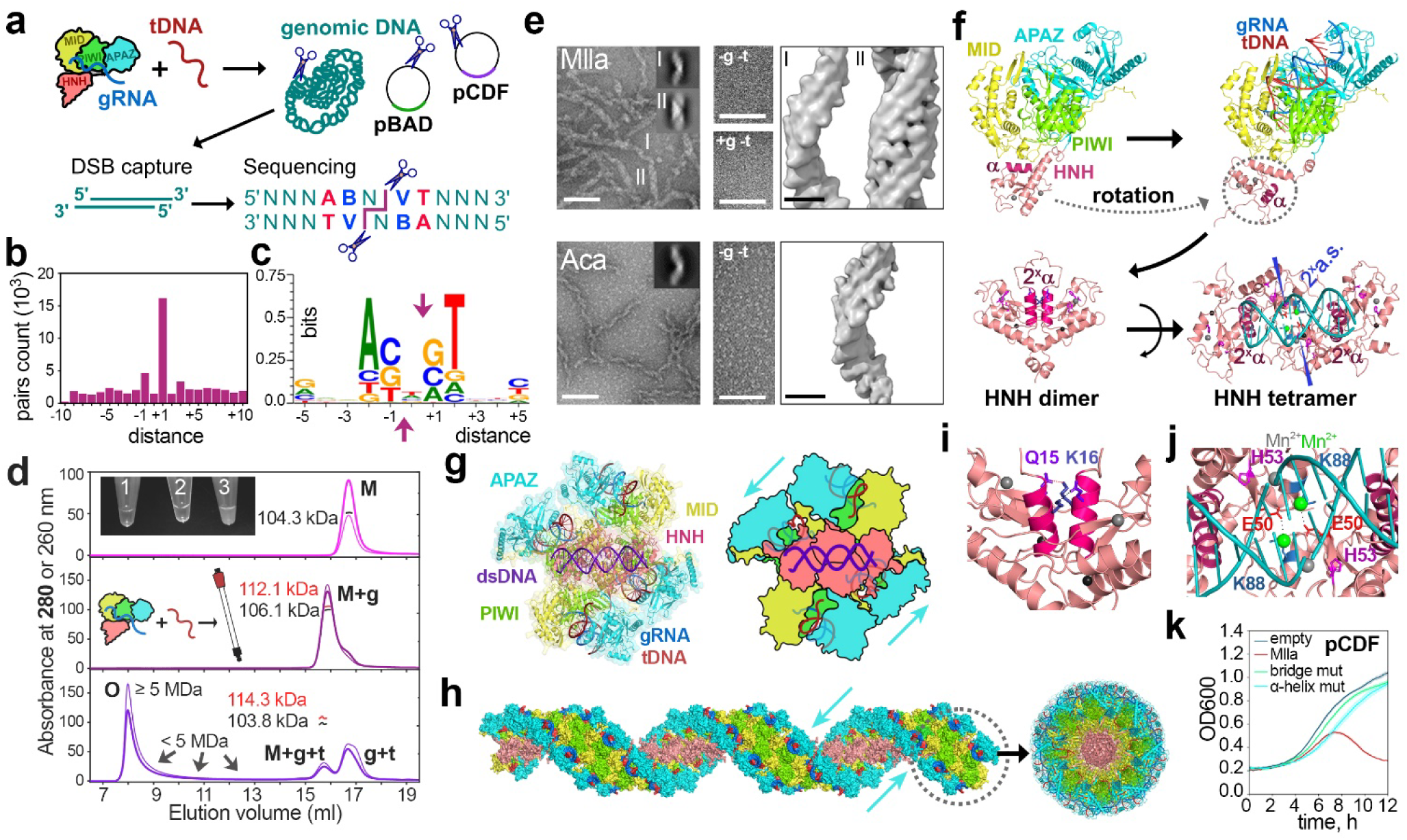
The mechanism of SPARHA activation. **a,** Mapping of the sites of DNA cleavage by SPARHA. **b,** Distribution of distances between closest cleavage sites on the opposite DNA strands for MllaSPARHA. **c,** Nucleotide logo of the consensus cleavage site of MllaSPARHA, calculated for double-cleavage sites with a one-nucleotide distance. **d,** Analysis of the oligomerization state of SPARHA (CD mutant of AcaSPARHA) by SEC-MALS, in the absence of guides/targets (top), in the presence of guide RNA (middle), or guide RNA and complementary target DNA (bottom). The profiles show absorbance at 280 (thick) and 260 (thin) nm. The measured MWs of nucleic-acid-free (104.3 kDa), guide-bound (112.1 kDa RNA-protein, 106.1 kDa protein) and guide-target-bound (114.3 kDa RNA-DNA-protein, 103.8 kDa protein) SPARHA are indicated. M, monomers; O, oligomers of heterodimeric SPARHA. The inset shows samples of CD AcaSPARHA containing non-complementary (1 and 2) or complementary (3) guide-target combinations. **e,** TEM images of MllaSPARHA and AcaSPARHA in the presence of complementary guide RNA and target DNA (left; 2D class averages are shown in insets) and in the absence of guides and targets (middle), scale bar 100 nm. 3D density maps of filaments (right), scale bar 10 nm. **f,** Proposed pathway of SPARHA activation. The α helix involved in HNH dimerization is shown in dark pink. Catalytic HNH residues holding a Mn^2+^ ion are shown in magenta. The activated HNH tetramer containing a double active site (2^x^a.s.) is shown in complex with a dsDNA substrate. Structural metal ions at the interface between HNH dimers are shown in green. Positions of DNA cleavage are indicated with arrows. **g,** Model of a tetramer of heterodimeric MllaSPARHA with bound guide RNA, target DNA and substrate dsDNA. **h,** Modelled filament of activated MllaSPARHA, side and end views. **i,** Interactions between the N-terminal α helixes of HNH domains in the HNH dimer. The Q15/K16 residues involved in dimerization are shown. **j,** Interface between active monomers in the model of HNH oligomer with a dsDNA substrate; DNA cleavage sites are shown as breaks in the strands. H53 residues holding catalytic Mn^2+^ ions (gray) are shown in magenta. E50 and K88 residues from opposite HNH monomers holding structural metal ions (Mn^2+^, green) are shown in red. All models were generated using AF3, with manual adjustment of HNH positions in the tetramer of MllaSPARHA with substrate DNA, based on the experimental structure of Cap5. **k,** Growth of *E. coli* strains expressing WT MllaSPARHA or its mutants with alanine substitutions in the N-terminal α helix (Q15A/K16A) or at the active site interface (bridge mutant, E50A) in the presence of pCDF. Means and standard deviations from three independent experiments. See Fig. ED5g for cell grown in the absence of pCDF.

To reveal whether SPARHA complexes undergo oligomerization upon activation, we performed size-exclusion chromatography coupled to multi-angle light scattering (SEC-MALS) of apo- and nucleic-acid-bound SPARHA. Apo-SPARHA was a heterodimeric complex of pAgo and HNH-APAZ of the expected molecular weight (Fig. 4d, Fig. ED9a). Target DNA was not bound by SPARHA in the absence of guide RNA (Fig. ED9b). Guide RNA was bound by SPARHA without formation of higher-order structures (Fig. 4d, Fig. ED9b). Further addition of target DNA to the guide-SPARHA complex resulted in instantaneous oligomerization of SPARHA, with most nucleic-acid-bound complexes eluting in the column void volume (apparent MW of >5 MDa, corresponding to >45 heterodimeric SPARHA complexes), a shoulder of smaller oligomeric complexes with MW <5 MDa, and a small peak of heterodimeric guide-target-bound SPARHA (Fig. 4d, Fig. ED9b,c). The oligomerization of SPARHA resulted in visible changes in the transparency of the solution; the samples became opaque in the presence of guide RNA and complementary target DNA, but not without target DNA or guide RNA, or with noncognate target DNA or guide RNA (Fig. 4d inset, Fig. ED9d).

The activated SPARHA complexes were visualized by negative staining transmission electron microscopy (TEM). Both MllaSPARHA and AcaSPARHA formed highly regular corkscrew-like filaments with a characteristic length of several hundred nanometers and an average helix diameter of 16 nm (Fig. 4e, Fig. ED10a,b). For MIlaSPARHA, an additional filament type was observed, characterized by a distinct structure that maintains the same diameter but appears straight rather than corkscrew-like (Fig. 4e). No higher-order structures were observed in the absence of guides and targets or in the presence of guide RNA only (Fig. 4e, Fig. ED10a,b). 2D classification of filament segment projections, followed by 3D reconstruction of the helical segment density map, revealed helical filaments for both MIlaSPARHA and AcaSPARHA, with distinct ‘vertebrae’ structures, likely corresponding to individual SPARHA complexes (Fig. 4e, Fig. ED10a,b). The estimated helical parameters are a helical rise of 52 Å and a helical twist of 38°. The second type of MIlaSPARHA filament was identified as a double helix consisting of two intertwining corkscrew-like helices (Fig. 4e, Fig. ED10a).

Based on analysis of DNA cleavage by SPARHA, SEC, TEM and structural modelling, we propose the following scenario of target-induced activation of SPARHA (Fig. 4f). Activation of the Cap5 effector, containing a closely related HNH domain (Fig. 1f), involves tetramerization of HNH, with a double active site formed at the interface between two HNH dimers (Fig. ED11d) ^27^. Individual HNH dimers in Cap5 are stabilized by interactions between N-terminal α helixes, while the active site interface between two dimers is stabilized by additional non-catalytic metal ions interacting with glutamate residues from each dimer, present in both Cap5 and SPARHA (Fig. 4f, Fig. ED11d). AF3 modelling shows that the N-terminal helix of HNH in apo-SPARHA interacts with the MID domain of pAgo, close to the guide-binding pocket of MID, likely corresponding to the inhibited state of HNH. The binding of guide RNA and target DNA to SPARHA likely induces HNH rotation, freeing its N-terminal helix and making possible dimer formation (Fig. 4f, Fig. ED11a). Further dimerization of HNH dimers generates an active HNH tetramer with a double active site formed in between central HNH monomers and containing a pair of structural metal ions (Fig. 4f), similarly to Cap5 (Fig. ED11c,e). The DNA substrate can be perfectly fit into the catalytic cavity, allowing cleavage of the two strands at a one-nucleotide distance (Fig. 4f,g). Additional dimers of the heterodimeric SPARHA complex can likely attach to the tetramer in a head-to-tail orientation to extend the filament at both sides (Fig. 4h). In the resulting filament model, the pAgo/APAZ blocks form a right-handed spiral scaffold that can potentially hold a continuous stretch of active HNH domains in its center (Fig. 4h).

The proposed structural model fit well into the 3D density map derived from TEM images (Fig. 4e, Fig. ED10a,b) and was verified by analysis of SPARHA mutants. Substitutions of glutamine and lysine residues in the N-terminal helix of HNH involved in dimer formation (Q15A/K16A, Fig. 4i) fully relieved HNH toxicity in the presence of the invader plasmid (Fig. 4k, α helix mutant). Substitution of the glutamate residue (E50A) coordinating the structural metal ion at the active interface between HNH dimers (Fig. 4j, green) similarly inactivated SPARHA (Fig. 4k, bridge mutant). This confirmed that both types of interactions between HNH domains in the HNH filament are crucial for the nuclease activity of SPARHA. Further high-resolution structural analysis will be required to confirm the proposed model and to reveal the structure of the second type of filaments formed by MllaSPARHA.

## Discussion

SPARHA is a inducible immune system with an HNH effector that is held inactive in the absence of target DNA. Recognition of invader elements by SPARHA induces a cascade of architectural changes leading to the formation of HNH filaments and indiscriminate collateral degradation of cellular DNA (Fig. 5a). This likely protects bacterial population through an abortive infection mechanism, also employed by several other defense systems containing DNase effectors (STAND/NACHT, Hachiman, CBASS, Type III CRISPR) ^27,29-32^. In comparison with these systems, SPARHA directly converts guide-dependent sensing of specific invader DNA by pAgo into the nonspecific activity of ‘megalonuclease’ filaments. Given that short pAgos might have been an ancestor form of Argonautes ^3,5^, the evolution of Argonaute-based immunity likely went from supramolecular complexes that elicit indiscriminate suicide response (NADases and nucleases in short pAgo systems, Fig. 1a) to long eAgos specifically acting on individual mRNA targets.

**Figure 5.**
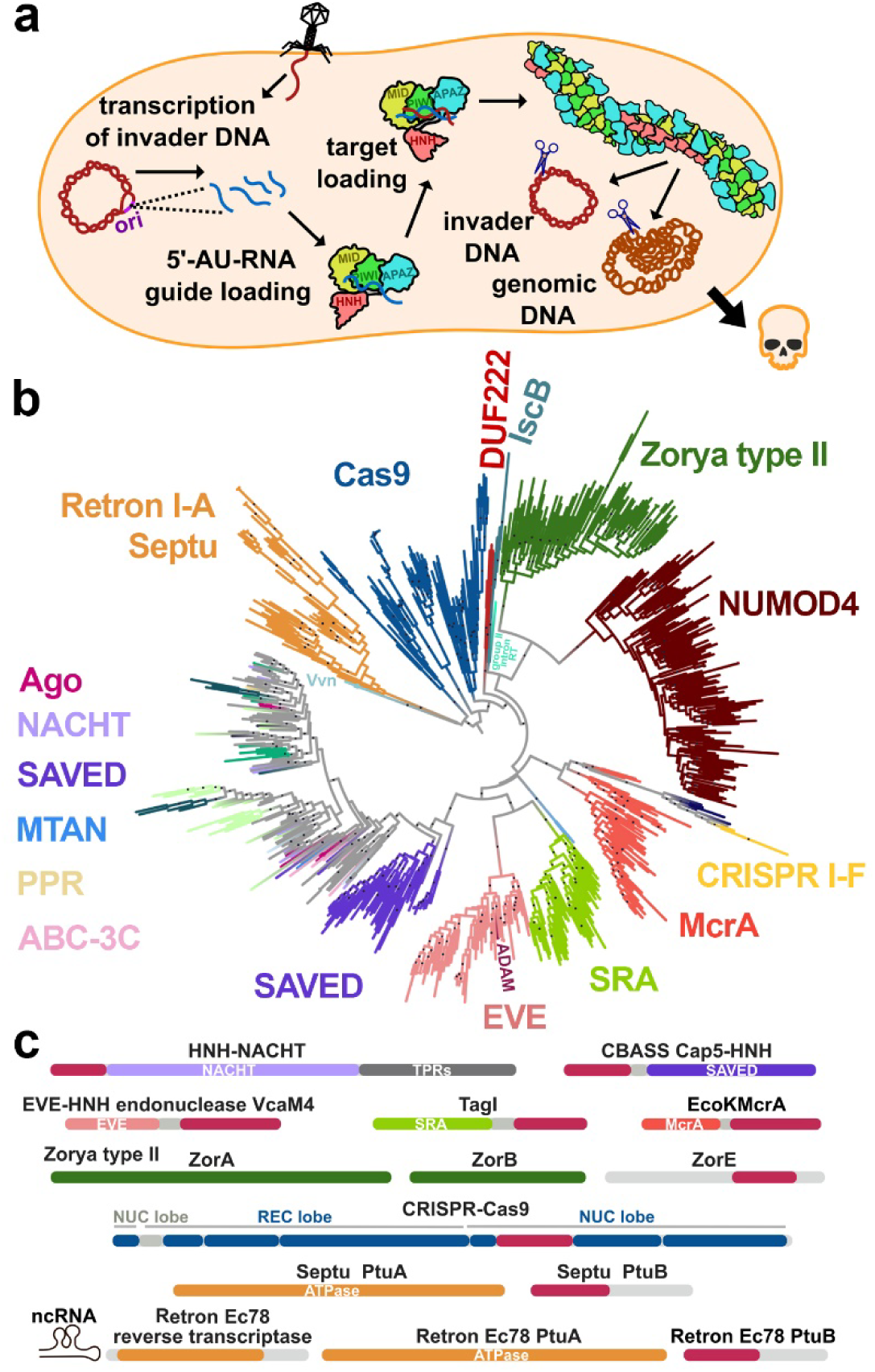
The role of SPARHA and HNH nucleases in immunity. **a,** Proposed mechanism of SPARHA. **b,** Phylogenetic tree of HNH-containing proteins, based on the sequence alignment of HNH domains. The dots represent bootstrap values from 85% to 100%. For each clade, the most abundant effector type is indicated. **c,** Representative defense systems containing HNH effectors. Panel **a** created with Inkscape (http://www.inkscape.org).

The observed preference of SPARHA for guide RNAs generated from the plasmid replication origin suggests that SPARHA may recognize specific features of invader elements. Certain preferences for plasmid transcripts were also reported for the SPARSA, SPARTA and SPARDA systems ^19,22,24^, but only for SPARHA plasmid-derived guide RNAs become dominant among other guide species. RNA-based origins of replication depend on host RNA polymerase for primer RNA synthesis and are commonly found in single-stranded DNA phages and conjugative plasmids. Initially transcribed regions of such plasmids are hotspots of various anti-defense factors, indicating that they are targeted by multiple immune systems ^33^, likely including short pAgo systems. High levels of transcription and high copy numbers of invader plasmids and phages may further induce immune response, as previously proposed for some CRISPR and pAgo systems ^19,24,34,35^. The requirement for filament formation during activation of SPARHA may act as a safeguard mechanism against self-immunity, by requiring high loads of specific RNA guides and invader DNA targets for the effector complex assembly.

HNH nucleases are widespread in prokaryotic defense systems, where they act as single domains and nick single-stranded DNA or form dimers to cleave both DNA strands, often within larger effector complexes (Fig. 5b,c). Monomeric HNH domains of Cas9 and its predecessor IscB, which diverge significantly from HNH of SPARHA (Fig. 5c and Fig. ED11g), are arguably the best known examples of HNH nucleases, acting in type II CRISPR immunity ^36^. Cas8-HNH effectors are also embedded in the Cascade complex of rare type I-F systems ^37^. Monomeric Cas9-related HNH domains are present in Septu and retron (Ec7/Ec78) effectors ^38,39^ and in Zorya type II systems ^40-42^, which introduce single-strand nicks in their DNA targets. Related HNH domains are also found in a large group of homing endonucleases, including monomeric nicking enzymes (fused to a NUMOD4 DNA-binding domain ^43^). In contrast, HNH domains of many modification-dependent restriction endonucleases act as dimers, recruited to their DNA substrates by various types of methylated DNA-binding domains (McrA, SRA, EVE or ADAM/DUF3578) ^44-48^. Finally, HNH effectors found in CBASS and SPARHA systems form oligomers upon activation (Ref. ^27^ and this study). The Cap5 effector of CBASS is activated by a cyclic dinucleotide binding to its SAVED domain, leading to HNH tetramerization ^27^; other HNH-SAVED effectors presumably employ similar mechanisms of activation. SPARHA systems are activated by the binding of guide-target to pAgo, showing that HNH effectors can be similarly controlled by various sensor/regulatory platforms. Remarkably, HNH nucleases from the diverse clade including SPARHA systems are found in association with a large variety of other domains with potential roles in immunity (Fig. 5c), including STAND ATPases (NACHT modules fused to tetratricopeptide repeat domains, TPR, and other sensor domains) ^32,49^, RNA-binding pentatricopeptide repeat domains (PPR) ^50^, effectors of ABC-3C ATPase systems ^51^ and MTAN nucleosidases ^52^ (Fig. 5c). We hypothesize that these systems also form tetramers or higher-order oligomers upon activation.

Oligomerization has emerged as a common theme in the action of prokaryotic immune systems ^53,54^. The binding of cyclic dinucleotides to the sensor domains induces formation of activated TIR filaments (TIR-SAVED or TIR-STING) ^55,56^ or transmembrane Cap15 filaments^57^ in CBASS systems. Similarly, SIR2 filaments are formed during activation of Thoeris (SLOG-SIR2) ^58^ and STAND/NACHT (SIR2-STAND) ^59^ systems, and Lon-SAVED protease filaments are formed in CRISPR Type III systems ^60,61^. SPARHA provides the first example of pAgo-activated filamentous nuclease. The SgrAI restriction endonuclease forms short filaments after recognizing unmethylated target sites, thus enabling biased cleavage of invader DNA ^62^. In contrast, formation of SPARHA filaments likely ensures highly efficient degradation of cellular DNA when sensing infection. Trigger-dependent collateral nuclease activity of such systems can be used for sensitive DNA detection or targeted elimination of genetic elements and cell types *in vivo*.

## Extended Data Figures

**Figure ED1.**
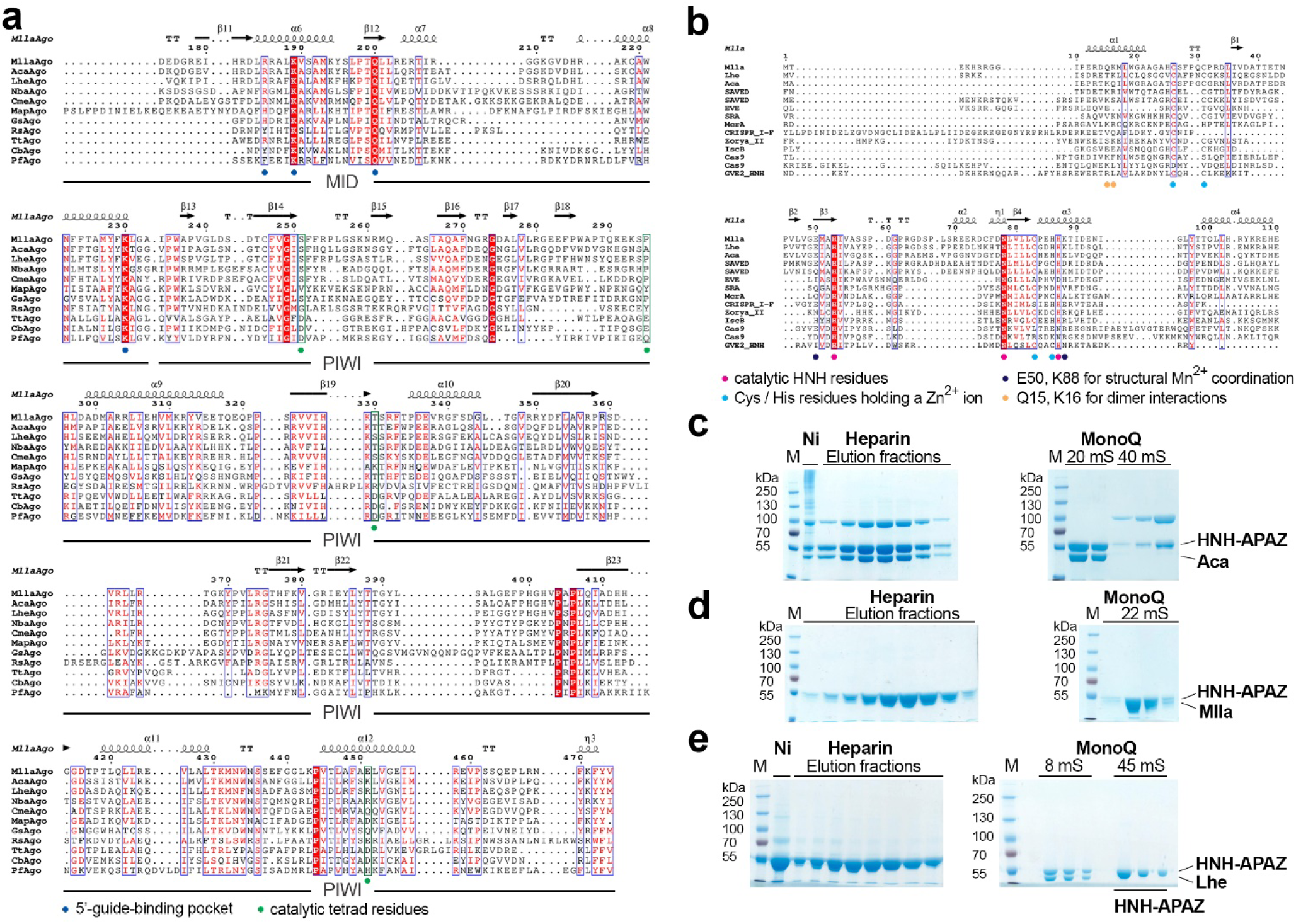
Sequence alignments of SPARHA systems and purification of SPARHA. **a,** Alignment of the MID and PIWI domains in short pAgos from SPARHA (MllaAgo, AcaAgo, LheAgo), SPARDA (NbaAgo, CmeAgo), SPARTA (MapAgo), SPARSA (GsAgo) systems and in long pAgos (RsAgo, TtAgo, CbAgo, PfAgo). Residues involved in interactions with the guide 5’-end in the MID pocket are shown with blue dots. The active site residues in the PIWI domain in active long pAgo nucleases (TtAgo, CbAgo, PfAgo) are shown with green dots. The source of pAgo proteins: MllaAgo – *M. llanfairpwllgwyngyllgogerychwyrndrobwllllantysiliogogogochensis*(WP_141644552.1), AcaAgo – *Acidithiobacillus caldus* (WP_215874059.1), LheAgo – *Leptolyngbya sp. ‘hensonii’* (WP_075600998.1), NbaAgo – *N. baekryungensis* (WP_022673743.1), CmeAgo – *C. metallidurans* (WP_011516870.1), MapAgo – *Maribacter polysiphoniae* (WP_109649955.1), GsAgo – *Geobacter sulfurreducens* (WP_010942012.1), RsAgo – *Cereibacter sphaeroides* (A4WYU7.1), TtAgo – *Thermus thermophilus* (WP_011174533.1), CbAgo – *Clostridium butyricum* (WP_058142162.1), PfAgo – *Pyrococcus furiosus* (WP_011011654.1). **b,** Residues of the HNH domain from various systems (from top to bottom): Mlla – SPARHA from *M. llanfairpwllgwyngyllgogerychwyrndrobwllllantysiliogogogochensis* (WP_141644551.1), Lhe – *Leptolyngbya sp. ‘hensonii’* (WP_075600999.1), SPARHA from Aca – *Acidithiobacillus caldus* (WP_215874058.1), SAVED – SAVED-HNH from *Pseudomonas sp. FSL W5-0299* (WP_077748502.1), SAVED – SAVED-HNH from *Lactococcus lactis* (WP_289448330.1), EVE – EVE-HNH from *Vibrio campbellii* (WP_010645282.1), SRA – SRA-HNH from *Thermocrispum agreste* (WP_084609162.1), McrA – McrA-HNH from *Streptomyces* (WP_011029780.1), CRISPR I-F – Cas8-HNH from *Selenomonas sp.* (WP_037363842.1), Zorya II – ZorE from *Escherichia coli* ATCC 8739 (ACA79492.1), IscB – (WP_235220879.1), Cas9 – *Geobacillus stearothermophilus* (WP_328074567.1), Cas9 – *Streptococcus pyogenes* (WP_373957964.1), GVE2 HNH – *Geobacillus virus E2* (YP_001285849.1). Catalytic HNH residues (magenta), cysteine/histidine residues holding a Zn^2+^ ion (cyan), residues coordinating non-catalytic Mn^2+^ ions (violet), and residues involved in interactions between HNH dimers in SPARHA (yellow) are indicated with dots. The amino acid numbering and secondary structure elements in panels a and b are shown for MllaAgo above the alignment (α-helices, π-helices and β-strands; strict β-turns are shown as TT). The alignment was created with ESPript 3.0. **c-e,** Purification of SPARHA complexes. **c,** Co-elution of AcaAgo and HNH-APAZ during Ni-chelating, heparin (left) and anion-exchange (MonoQ column, right) chromatography steps. **d-e,** The same for MllaAgo/HNH-APAZ and LheAgo/HNH-APAZ, respectively. Individual fractions and final protein samples are shown for the wild-type complex. In the case of LheAgo/HNH-APAZ, the fractions corresponding to HNH-APAZ after anion-exchange chromatography are underlined. The identity of proteins was confirmed by mass-spectrometry.

**Figure ED2.**
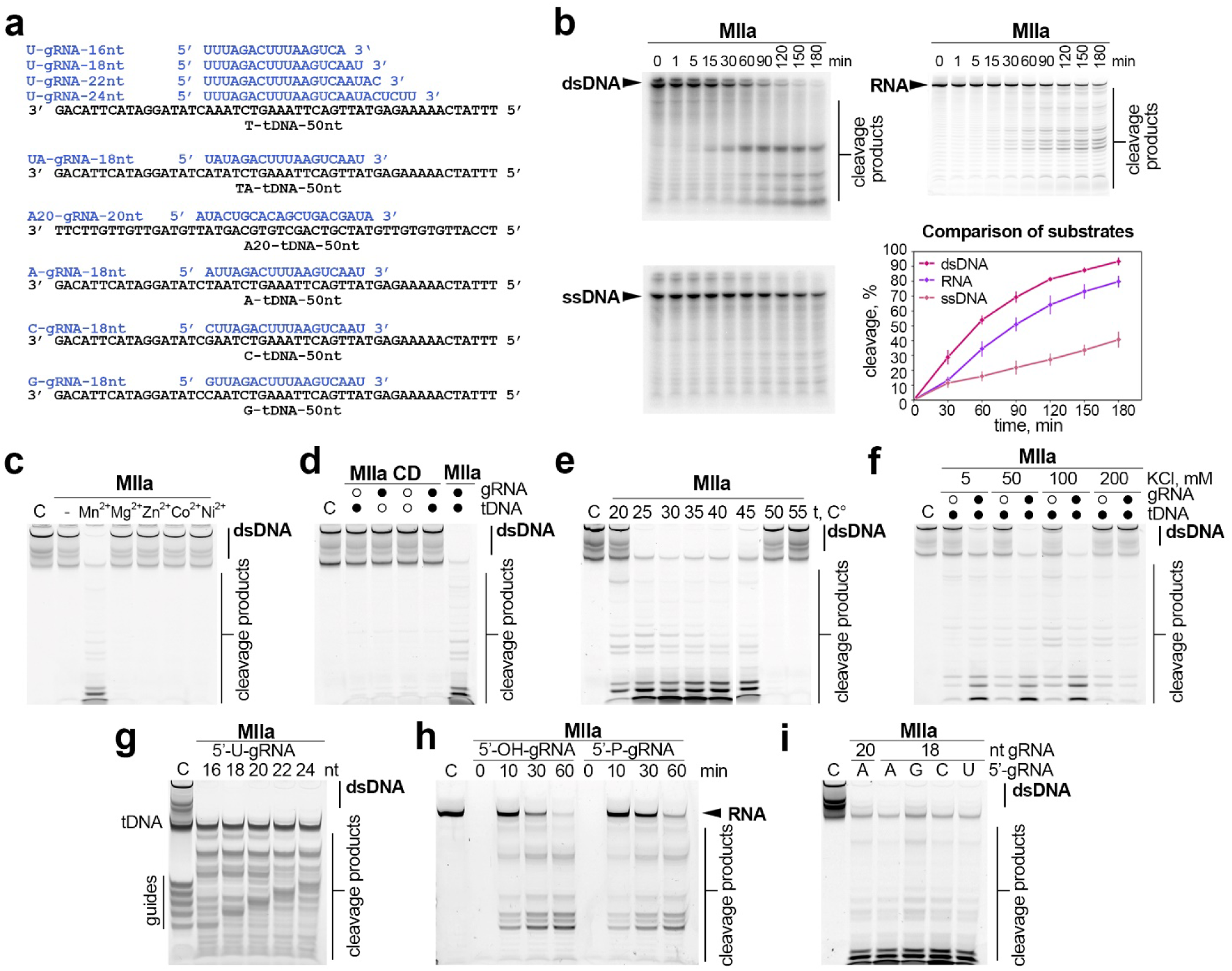
*In vitro* activities of MllaSPARHA. **a,** Sequences of guide RNAs (gRNAs) and target DNAs (tDNAs) used in the assays. **b,** Kinetics of cleavage of dsDNA, ssDNA and RNA substrates of identical sequences (55 nt long) by MllaSPARHA. The reactions contained 500 nM MllaSPARHA, 500 nM guide RNA and target DNA and 5 μM of collateral substrates. The DNA substrates were radiolabeled with ^32^P at the 5’-end. The RNA substrate contained a FAM dye at the 5’-end. The plot shows means and standard deviations from three independent experiments. **c,** Analysis of MllaSPARHA activity with various divalent cations (5 mM each, except for Zn^2+^ at 200 μM). **d,** Comparison of the activities of wild-type and catalytically dead (CD) MllaSPARHA. **e,** Temperature dependence of MllaSPARHA activity. **f,** Activity of MllaSPARHA in buffers with different ionic strength. **g,** Activity of MllaSPARHA with guide RNAs of various lengths. **h,** Comparison of the kinetics of RNA cleavage with 5’-P or 5’-OH guide RNAs. **i,** Comparison of the MllaSPARHA activity with guide RNAs containing various 5’-nucleotides. All reactions were performed at 30 °C in a reaction buffer containing 5 mM Mn^2+^, 100 mM KCl, pH 8.0 unless otherwise indicated. The reaction products were visualized by phosphorimaging (panel **b**, ssDNA and dsDNA substrates) or fluorescence scanning (FAM – RNA substrate, panels **b**, **h**; HEX – dsDNA substrate panels **c-f**, **i**) or SYBR Gold staining (panel **g**). Representative gels from at least three independent experiments are shown in each case. Positions of guide RNA, target DNA, collateral substrates and the cleavage products are indicated.

**Figure ED3.**
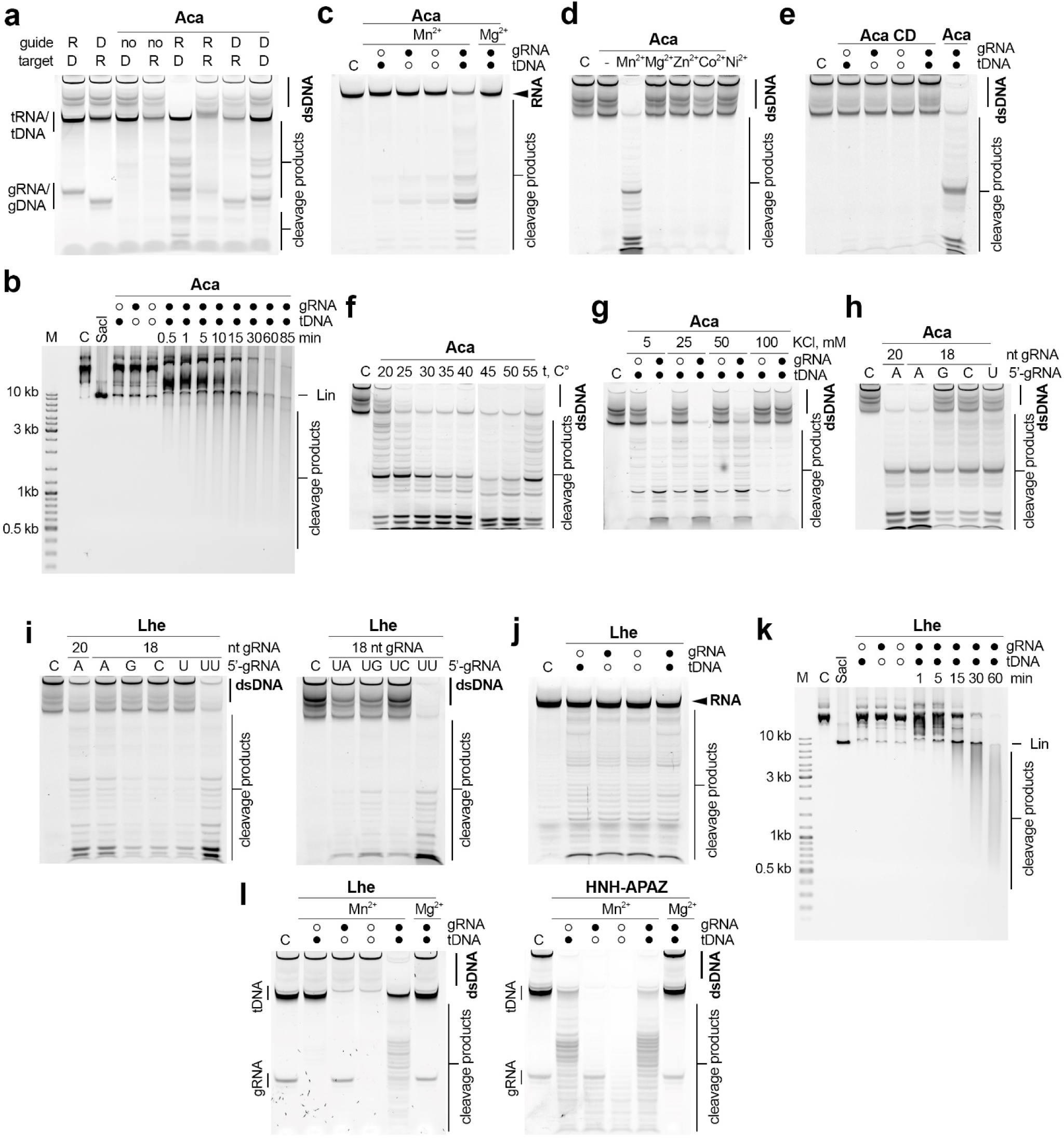
*In vitro* activities of AcaSPARHA (a-h) and LheSPARHA (i-l). **a,** Analysis of the specificity of AcaSPARHA for RNA (R) and DNA (D) guides and targets with a linear 55 nt dsDNA substrate. **b,** Kinetics of cleavage of plasmid DNA (pBAD) by AcaSPARHA in the presence of gRNA and tDNA. Linear plasmid (lane 3) was obtained by treatment with SacI. Control reactions were performed in the absence of gRNA (lane 4), or tDNA (lane 5), or both (lane 6). Lin, linear; M, length marker. **c,** Cleavage of 5’-FAM-labeled 55 nt RNA by AcaSPARHA. C, control without AcaSPARHA. **d,** Analysis of the AcaSPARHA activity with various divalent cations (5 mM each, except for Zn^2+^ at 200 μM). **e,** Comparison of the activities of wild-type and CD AcaSPARHA. **f,** Temperature dependence of the AcaSPARHA activity. **g,** Activity of AcaSPARHA in buffers with different ionic strength. **h,** Comparison of the activity of AcaSPARHA with guide RNAs containing various 5’-nucleotides. AcaSPARHA is efficiently activated only by gRNAs with 5’-A (lanes 2 and 3). **i,** (Left) Analysis of the activity of LheSPARHA with gRNAs containing various 5’-nucleotides. LheSPARHA is activated by gRNAs with 5’-UU and 5’-A (lanes 7 and 2). (Right) Effects of substitutions of the second nucleotide at the 5′-end in gRNA on the activity of LheSPARHA. The 5’-end dinucleotides of gRNAs are shown above the gel. C, control without AcaSPARHA. **j,** Activity of LheSPARHA with 5’-FAM-labeled 55 nt substrate RNA. **k,** Kinetics of cleavage of plasmid DNA (pBAD) by LheSPARHA in the presence of gRNA and tDNA. Linear plasmid (lane 3) was obtained by treatment with SacI. Control reactions were performed in the absence of gRNA (lane 4), or tDNA (lane 5), or both (lane 6). Lin, linear; M, length marker. **l,** Cleavage of the 55 nt dsDNA substrate by LheSPARHA (left) or LheHNH-APAZ (right). LheSPARHA degrades the dsDNA substrate only in the presence of gRNA and tDNA (lane 5). In contrast, LheHNH-APAZ degrades the collateral substrate independently of gRNA or tDNA (lanes 2-5) and also degrades tDNA when it is added to the reaction (lanes 2, 5). The gels correspond to Fig. 2g, but are visualized with fluorescence scanning. All reactions were performed at 30 °C in reaction buffers containing 5 mM Mn^2+^ (with 5 mM KCl for AcaSPARHA or 100 mM KCl for LheSPARHA), unless otherwise indicated. The reaction products were visualized by fluorescence scanning (HEX – panels **a, d-i**, **l**; FAM – panels **c**, **j**) or SYBR Gold staining (panels **b**, **k**). Representative gels from at least three independent experiments are shown in each case. Positions of gRNA, tDNA, collateral substrates and the cleavage products are indicated. See Table S2 for substrate sequences.

**Figure ED4.**
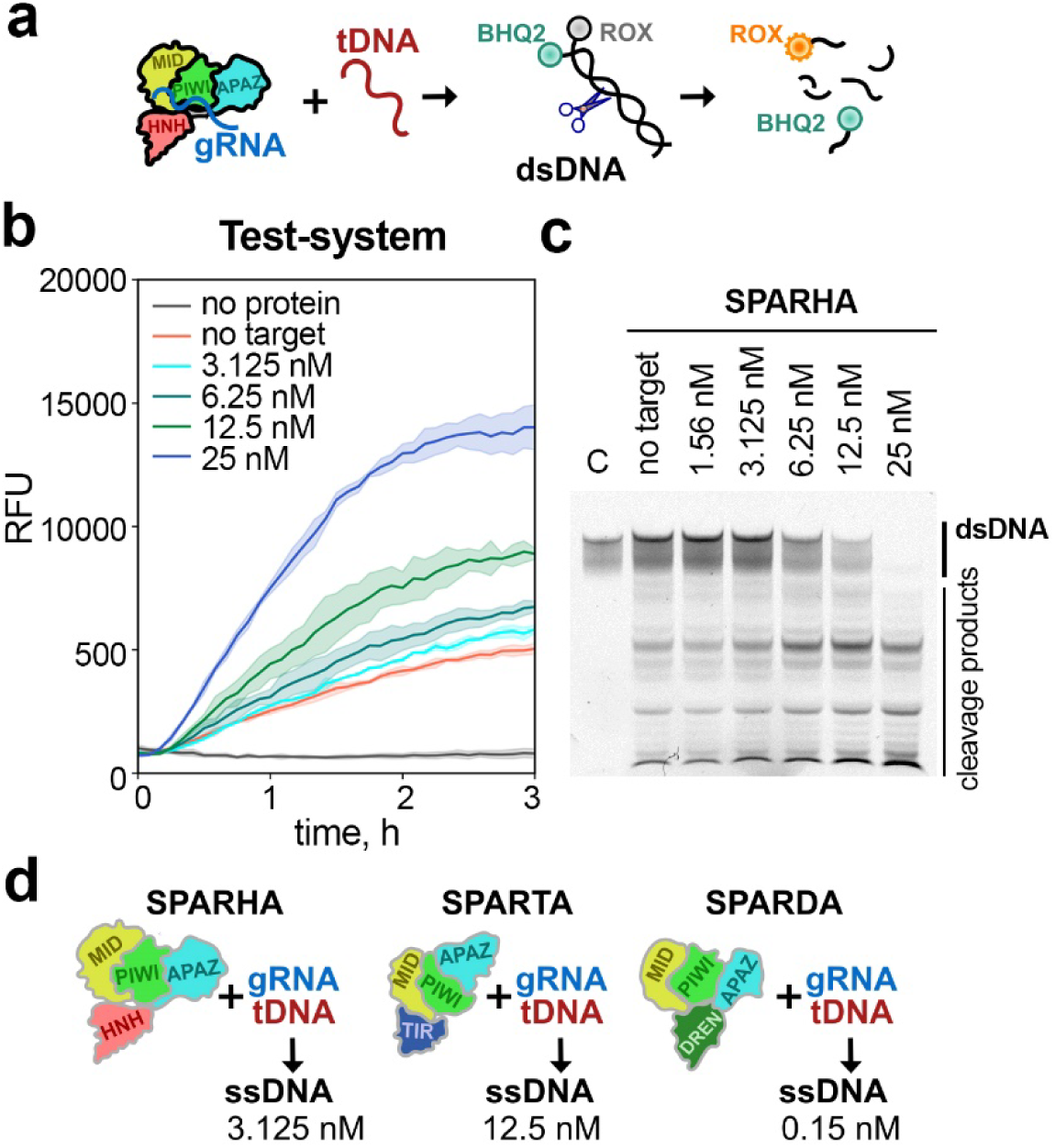
DNA detection with SPARHA. **a,** Scheme of the fluorescent beacon assay. **b,** Fluorescence readout after guide-dependent recognition of single-stranded target DNA (added to the final concentrations from 3.125 nM to 25 nM) by AcaSPARHA (1 μM, loaded with 1.3 μM gRNA), using an dsDNA beacon. Means and standard deviations from three replicates. RFU, relative fluorescence units. **c,** Visualization of the dsDNA beacon cleavage at 1.5 hours of the reaction by PAGE. The concentrations of target DNA are shown above the gel. The reaction products were visualized by fluorescence scanning (HEX). Position of the dsDNA beacon substrate and the cleavage products are indicated. See Table S2 for oligonucleotide sequences. **d,** Comparison of the reported sensitivities of test-systems based on the activity of short pAgos (SPARTA, SPARDA and SPARHA). Panels **a** and **d** created with Inkscape (http://www.inkscape.org).

**Figure ED5.**
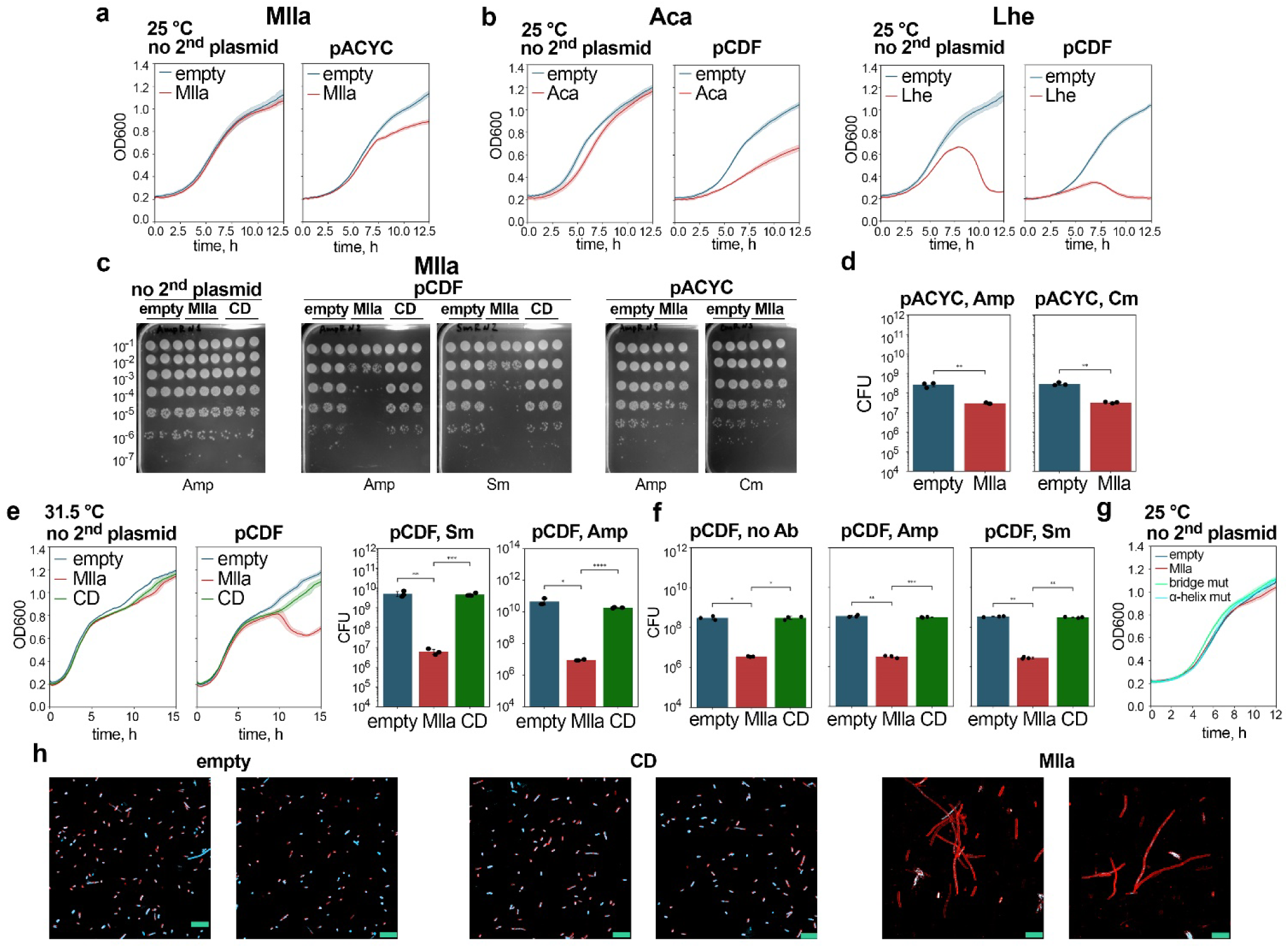
Analysis of the effects of MllaSPARHA on cell growth and viability in the presence of invader plasmids. **a,** Growth of *E. coli* strains expressing WT MllaSPARHA or containing empty pBAD, with or without the interfering plasmid pACYC, in the presence of Amp and 0.2% L-Ara (to induce SPARHA expression). Means and standard deviations from six independent experiments. **b,** The same for AcaSPARHA (left) and LheSPARHA (right) in the absence and in the presence of the interfering plasmid pCDF. **c,** Measurements of CFU numbers for *E. coli* strains containing pBAD encoding MllaSPARHA (WT or CD mutant) or empty pBAD (Amp^R^) in the absence (left) and in the presence of pCDF (Sm^R^, center) or pACYC (Cm^R^, right). The cells were grown for 8 hours at 25 °C (see Fig. 3b, right) in the presence of Amp and 0.2% Ara (to induce SPARHA expression), and serial dilutions were plated on LB agar containing Amp or Cm or Sm and 0.5% Glc (to repress SPARHA expression). CFU numbers were counted after overnight growth. **d,** Numbers of viable cells in cell cultures grown for 8 hours (shown in panel a), expressing MllaSPARHA and containing pACYC, measured in the presence of Amp or Cm. Means and standard deviations from 3 independent experiments. Statistically significant differences are indicated (from left to right: **p = 0.0044, **p = 0.0011; calculated using the one-sided t-test for independent samples). **e,** (Left) Growth of *E. coli* strains expressing WT or CD MllaSPARHA, or containing control empty pBAD, with or without the interfering plasmid pCDF at 31.5 °C in the presence of Amp and 0.2% L-Ara (to induce SPARHA expression). Means and standard deviations from six independent experiments. (Right) Numbers of viable cells in cell cultures grown for 10.5 hours (from the experiment shown on the left), expressing WT or CD mutant of MllaSPARHA and containing pCDF, measured in the presence of Amp or Sm. Means and standard deviations from 3 independent experiments. Statistically significant differences are indicated (from left to right: *p = 0.017, ***p = 0.00007, **p = 0.0076, ***p = 0.0007; calculated using the two-sided t-test for independent samples with Holm-Bonferroni correction). **f,** CFU numbers in cell cultures expressing WT or CD mutant MllaSPARHA or containing empty pBAD in the presence of pCDF, grown for 8 hours at 25°C in the presence of 0.2% L-Ara (to induce SPARHA expression) without antibiotics, and plated on LB agar without antibiotics or containing Amp or Sm, with the addition of 0.5% glucose (to repress SPARHA expression). Means and standard deviations from 3 independent experiments. Statistically significant differences are indicated (from left to right: *p = 0.015, *p = 0.01, **p = 0.0055, ***p = 0.0007, **p=0.0014, **p=0.0017; calculated using the two-sided t-test for independent samples with Holm-Bonferroni correction). **g,** Growth of *E. coli* strains expressing WT MllaSPARHA or its mutants, or containing empty pBAD, without interfering plasmids, in the presence of Amp and 0.2% L-Ara (to induce SPARHA expression). Corresponding growth curves of cell cultures in the presence of the interfering plasmid pCDF are shown in Fig. 4k. Means and standard deviations from three independent experiments. **h,** Staining of *E. coli* cells expressing WT or CD MllaSPARHA, or containing empty pBAD, in the presence of the interfering plasmid pCDF with DNA-specific (SYTO9, shown in cyan) and membrane-specific (SynaptoProbe Red) dyes. Cell cultures were collected at 7.5 hours of growth. The scale bar is 10 μm. Representative fields of view from three independent experiments (5–10 analyzed in each experiment) are shown.

**Figure ED6.**
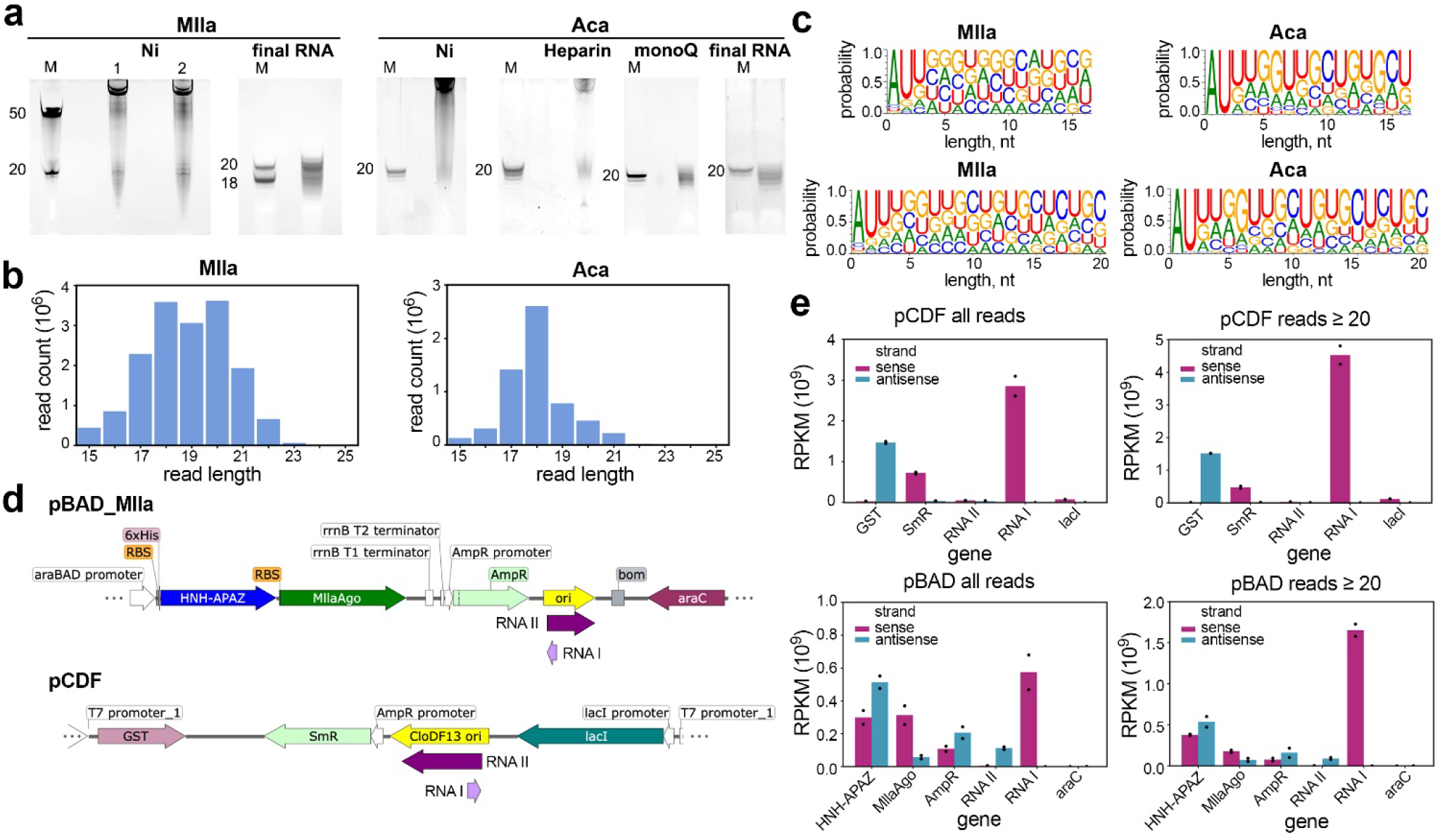
Analysis of guide RNAs bound to MllaSPARHA and AcaSPARHA in *E. coli*. **a,** Purification of small RNAs (∼15-25 nt) associated with MllaSPARHA (left) and AcaSPARHA (right) from *E. coli* containing the interfering plasmid pCDF. The samples were taken at different steps of chromatography (Ni-chelating, Heparin-affinity and MonoQ anion exchange) using a modified purification protocol with the exclusion of EDTA from buffers. The final fractions of small RNAs used for sequencing were taken after Ni-chelating chromatography for MllaSPARHA or after MonoQ chromatography for AcaSPARHA. M, length marker. Small RNAs were visualized by SYBR Gold staining. Representative gels from three independent experiments are shown. **b,** Length distribution of small RNAs associated with MllaSPARHA (left) and AcaSPARHA (right) determined after sequencing of RNA libraries. **c,** Sequence logos of gRNAs associated with MllaSPARHA (left) and AcaSPARHA (right) for all reads (top) and for reads ≥20 nt (bottom). **d,** Maps of the pBAD_MllaSPARHA (top) and pCDF (bottom) plasmids. **e,** Analysis of the distribution MllaSPARHA-associated gRNAs along plasmid sequences, for pCDF (top) and pBAD (bottom); all reads (left) and reads ≥20 nt (right). The amounts of short gRNAs are shown in RPKM (reads per kilobase per million mapped reads in the library) for each DNA strand (sense or antisense for each gene) and plotted for indicated plasmid regions. Means and standard deviations from two biological replicates.

**Figure ED7.**
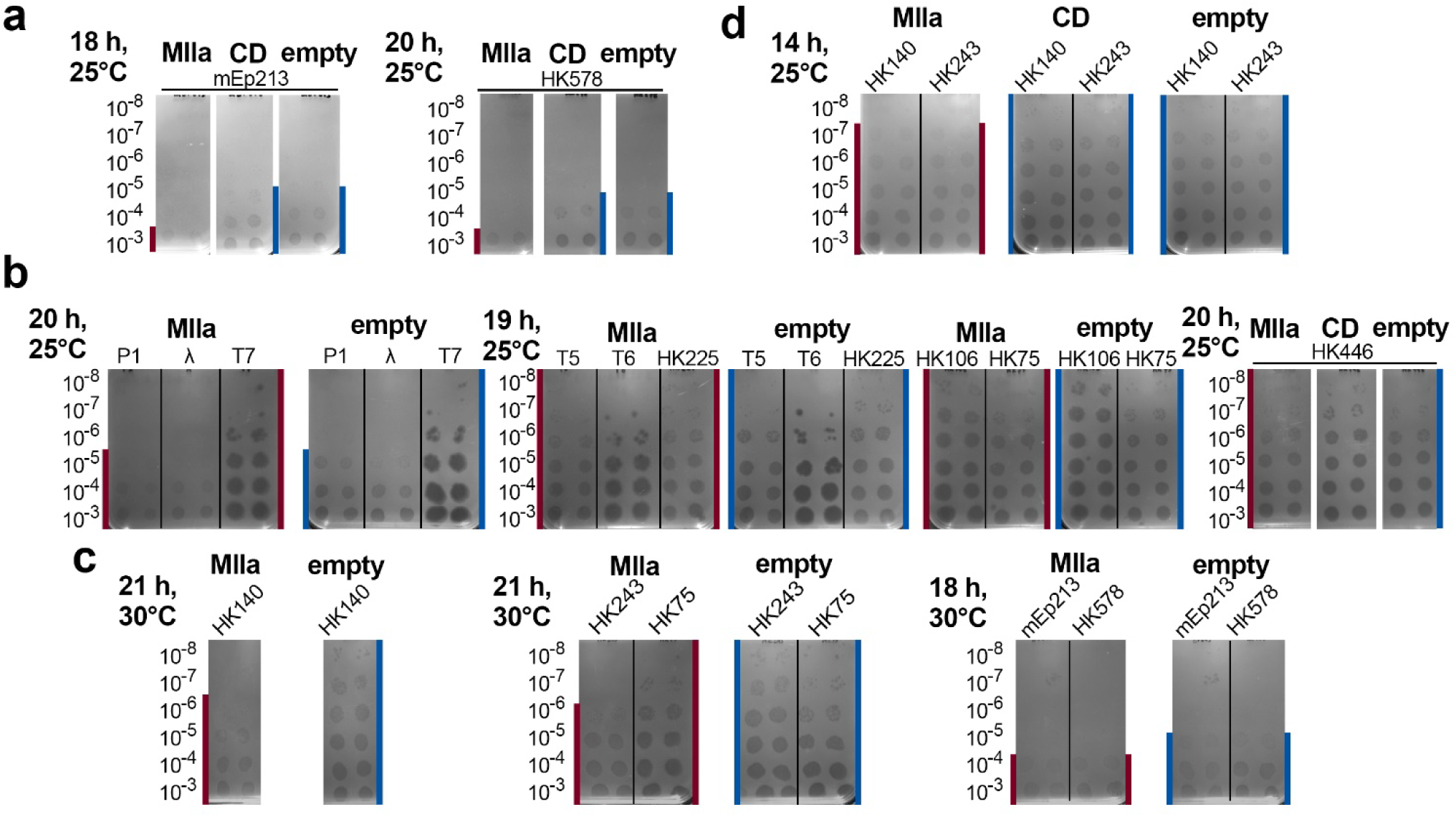
Analysis of the effects of MllaSPARHA on phage infection. **a,** Analysis of the EOP for phages mEp213 (left) and HK578 (right), which are sensitive to MllaSPARHA, with *E. coli* strains expressing WT or CD MllaSPARHA or containing empty pBAD. Representative plates from four independent experiments are shown (each containing two replicates). The corresponding EOP values are shown in Fig. 3i. **b,** Analysis of the EOP of the indicated phages with *E. coli* strains expressing WT MllaSARHA or containing empty pBAD (two replicates for each phage). In this panel, phages for which WT MllaSPARHA does not display any effects are shown. с, Analysis of the EOP of MllaSPARHA-sensitive phages HK140, HK243, HK75, mEp213, HK578 with *E. coli* strains expressing WT MllaSARHA or containing empty pBAD at 30 °C. Representative plates from four independent experiments are shown. **d,** Analysis of the EOP of phages HK140 and HK243 with *E. coli* strains expressing WT or CD MllaSARHA or containing empty pBAD, at 14 hours post-infection. The same plates with longer incubation time are shown in Fig. 3h. Serial dilutions of phage stocks and the experimental conditions are indicated on the left. The maximum dilutions with visible plaques or spots are indicated by vertical lines on each panel (red for WT MllaSPARHA, blue for CD MllaSPARHA and empty pBAD).

**Figure ED8.**
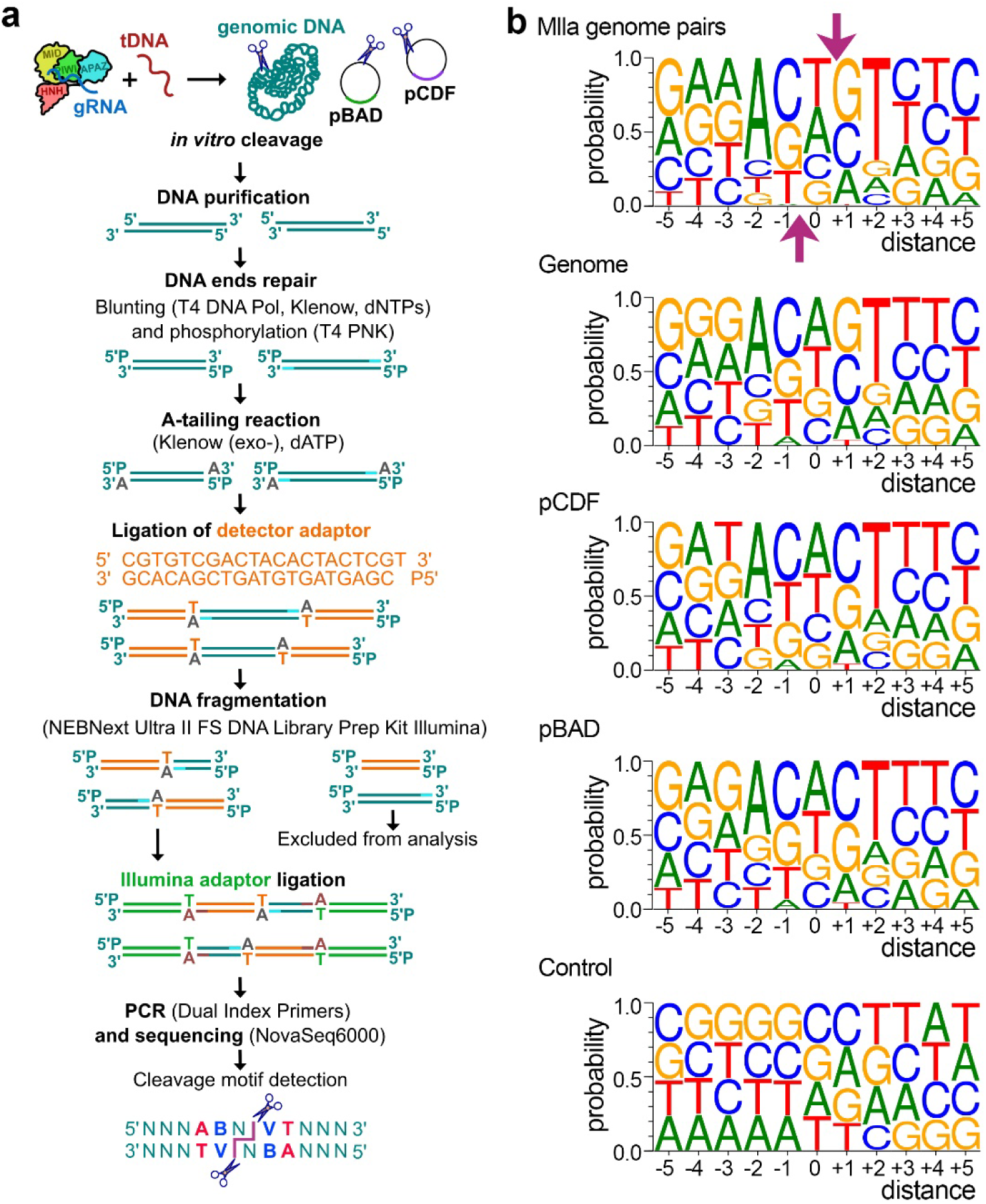
Mapping of the sites of DNA cleavage by MllaSPARHA. **a,** Protocol for DSB mapping. **b,** Sequence logos of DSBs in genomic and plasmid DNA for MllaSPARHA in comparison with the combined logo (genome and plasmids) for the control sample obtained without treatment by MllaSPARHA (bottom). The top logo shows nucleotide preferences for DSBs containing nicks separated by a one-nucleotide distance (corresponding to the +1 peak in Fig. 4b); positions of DNA cleavage on each strand are indicated with arrows. The remaining logos show nucleotide preferences for all identified sites of cleavage on one DNA strand.

**Figure ED9.**
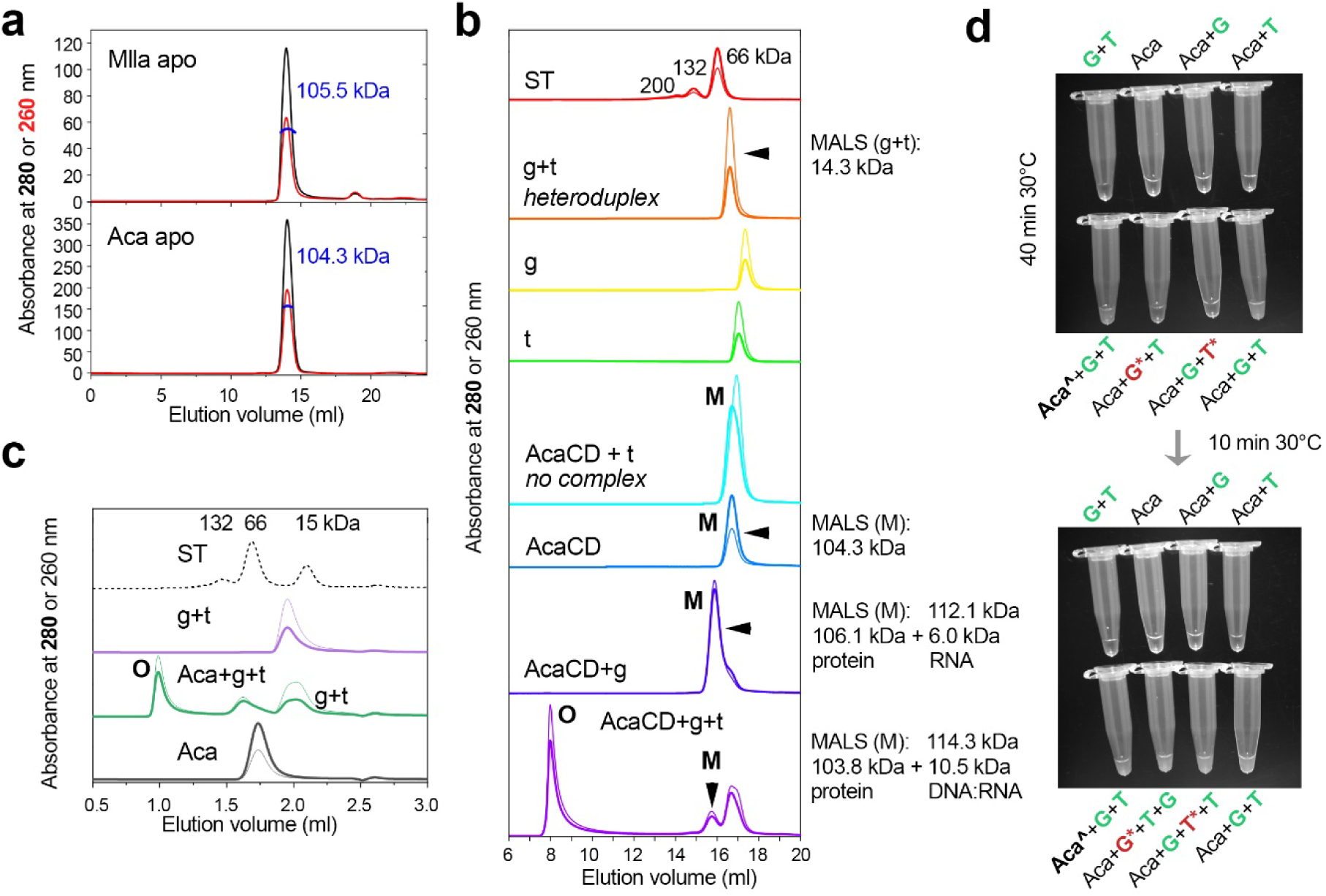
Analysis of the oligomeric state of SPARHA complexes. **a,** SEC-MALS profiles of heterodimeric complexes of MllaSPARHA and AcaSPARHA (at 280 nm, black, and 260 nm, red), using a Superdex 200 Increase 10/300 column. The calculated MWs of each complex are indicated. **b,** SEC-MALS analysis of CD AcaSPARHA, using a Superose 6 Increase 10/300 column. The profile containing BSA and α-lactalbumin standards (ST) is shown on the top, and their MW values are indicated in kDa (200 kDa corresponds to the BSA trimer). Peaks corresponding to monomers (M) and oligomers (O) of the heterodimeric SPARHA complex are indicated. The calculated MWs of the guide-target heteroduplex (g+t), heterodimer of AcaSPARHA (AcaCD), heterodimer in complex with guide RNA (AcaCD+g) and heterodimer in complex with guide RNA and target DNA (AcaCD+g+t) are indicated on the right of the plots, with contributions from the protein and nucleic acid parts; corresponding peaks are shown with arrowheads. **c,** SEC analysis of wild-type AcaSPARHA, using a Superdex 200 Increase 5/150 column. **d,** Photographic images of CD AcaSPARHA solutions, in the absence and in the presence of guides and targets. Aca CD was present at 5 μM concentration (3.3 μM in the sample labeled Aca CD^), the concentrations of all gRNA and tDNA were 3.3 μM. Guide RNAs and target DNAs were added as indicated. Complementary gRNA and tDNA are shown in green; non-complementary are shown in red with an asterisk. The photograph was taken after incubating the samples for 40 minutes at 30 °C. In all cases, the sample volume was 30 μL in a buffer containing 50 mM KCl (pH 7.0) with 5 mM Mn^2+^. The turbidity of the solutions is greatly increased only in the samples where complementary gRNA and tDNA were added. In the bottom photograph, the same samples are shown, except that correct gRNA/tDNA (3.3 μM) were added to the samples containing noncomplementary gRNA/tDNA pairs, which increased their turbidity (incubation for 10 minutes at 30 °C).

**Figure ED10.**
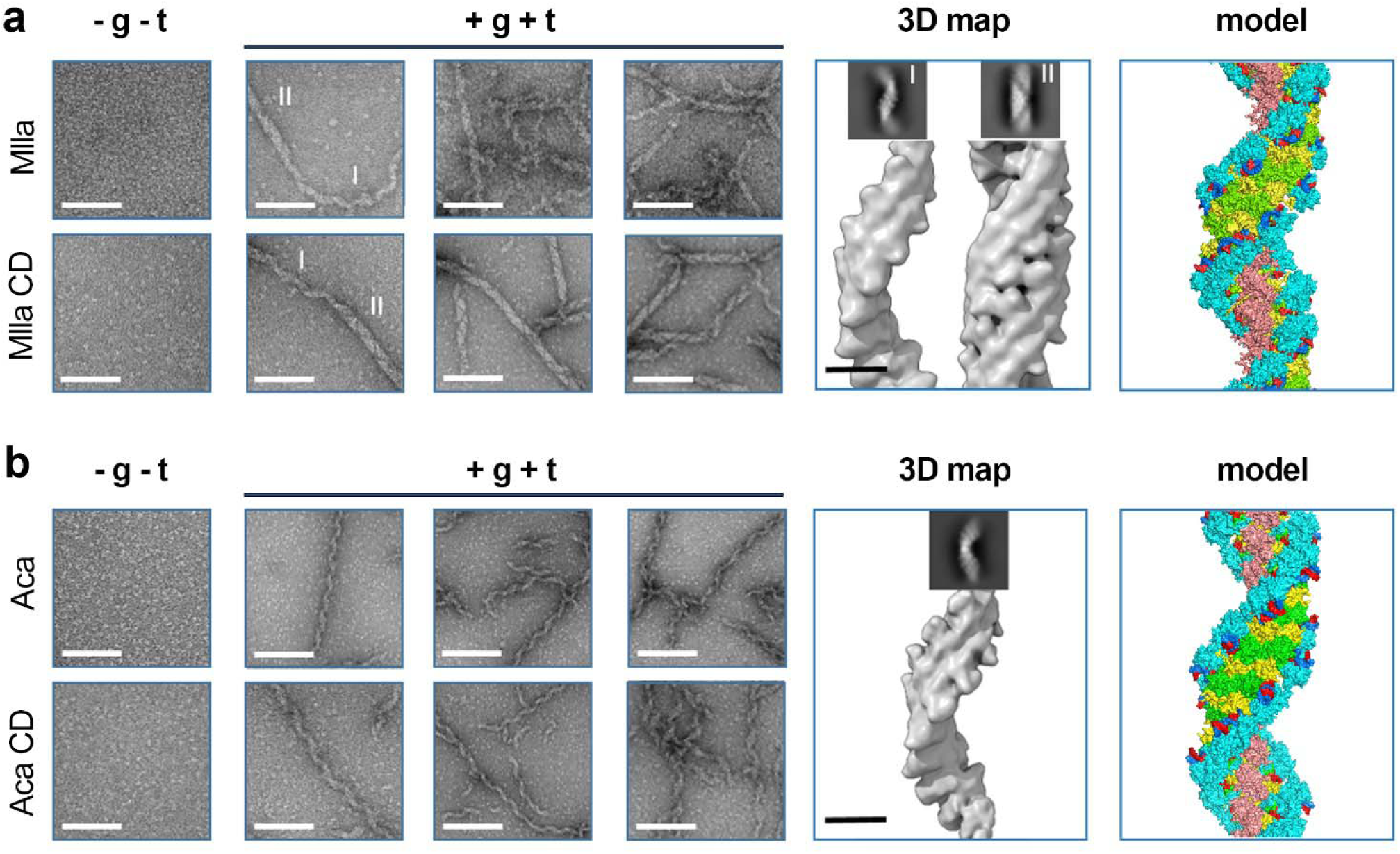
TEM images of MllaSPARHA (a) and AcaSPARHA (b). The samples were prepared with wild-type and CD SPARHA complexes. (Left) Negative-staining TEM images showing control samples without guides and targets and three frames of SPARHA filaments formed in the presence of guide RNA and target DNA. Scale bar: 100 nm. (Center) Corresponding 2D class averages displaying the averaged projections of individual filament segments and 3D reconstructions; scale bar: 10 nm. The 3D reconstruction (density map) of the filaments reveals that the filaments of MllaSPARHA exist in two distinct forms: single-helix and double-helix, whereas filaments of AcaSPARHA are exclusively single-helical. (Right) Proposed atomic models of the filaments created with UCSF ChimeraX based on AF3 predictions.

**Figure ED11.**
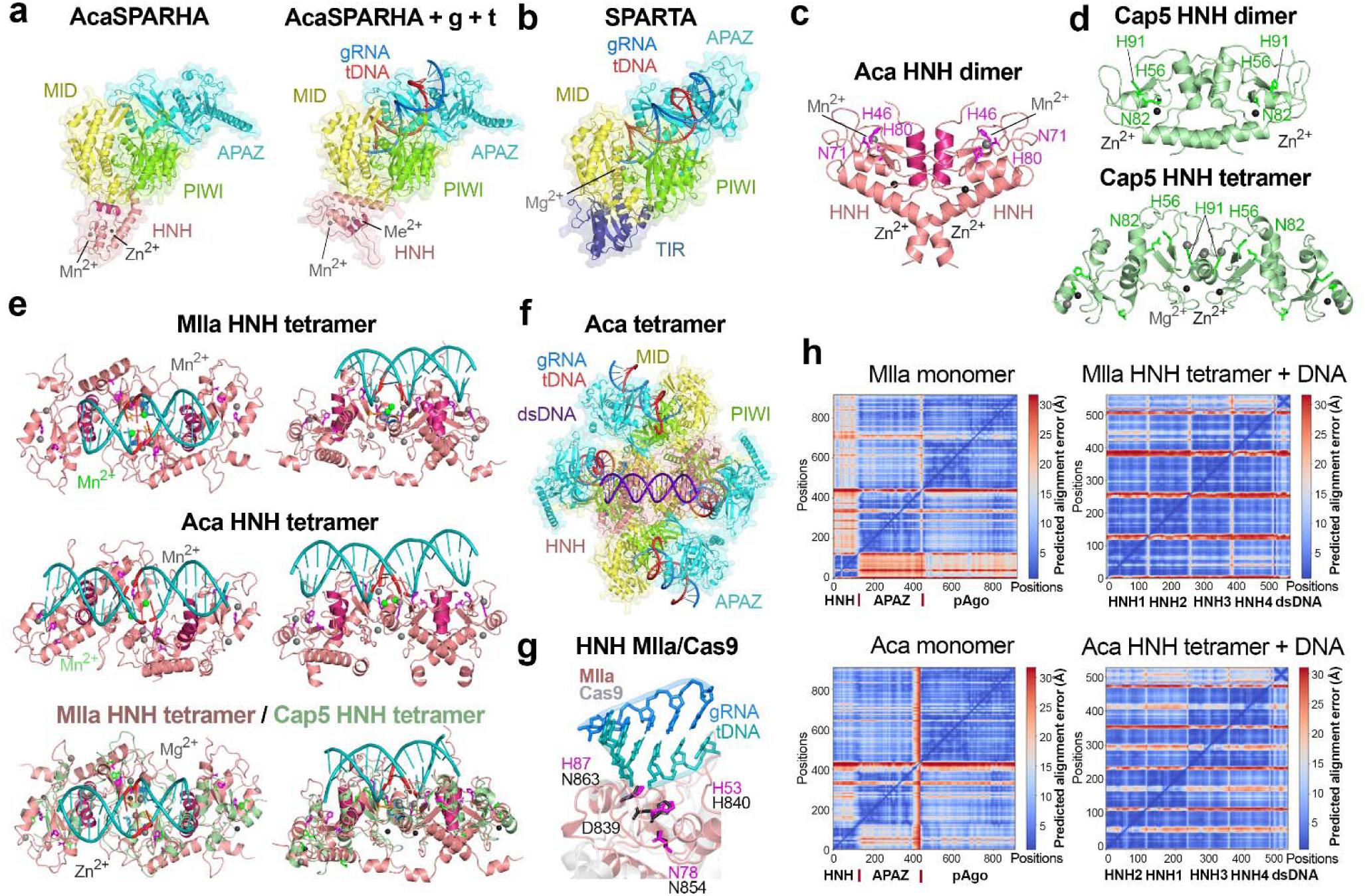
AlphaFold 3 models of MllaSPARHA and AcaSPARHA. **a,** AF3 models of the heterodimer of AcaSPARHA complex (left) and its complex with gRNA and tDNA (right). The APAZ domain is shown in cyan, the MID domain in yellow, the PIWI domain in green, and the HNH domain in pink. The N-terminal α-helix in the HNH domain involved in dimerization of HNH domains is shown in dark pink. **b,** The structure of the SPARTA system containing bound to guide RNA and target DNA (PDB ID: 8FFI). **c,** AF3 model of the Aca HNH dimer. The catalytic residues are shown in magenta. The α-helices that stabilize the dimer are shown in dark pink. **d,** The structure of the dimer (top, PDB ID:8FM1) and tetramer (bottom, PDB ID:8FMF) of the HNH domain of Cap5. The catalytic residues are shown in light green. **e,** AF3 models of DNA-bound tetramers of MllaSPARHA (top) and AcaSPARHA (middle) in two orientations, and structural alignments (bottom) of the DNA-bound tetramer of Mlla HNH (pink) with the HNH tetramer of Cap5 (green, PDB ID:8FMF). The DNA-bound model of the Mlla HNH tetramer was obtained by manual adjustment of the AF3 model, based on the Cap5 structure. The catalytic residues are shown in magenta; the dsDNA substrate is cyan, with the cleavage sites indicated in red. The α-helices stabilizing the dimers are shown in dark pink. **f,** AF3 model of a tetrameric complex of AcaSPARHA containing gRNAs, tDNAs, and dsDNA substrate. **g,** Structural alignment of the AF3 model of the Mlla HNH domain (pink) with the HNH domain of Cas9 (gray) containing sgRNA (blue) and target DNA (teal) (PDB ID: 6O0Y). The catalytic residues are shown in magenta (Mlla HNH) and black (Cas9 HNH). **h,** PAE (Predicted Aligned Error) maps for AF3 models of monomeric heterodimers of MllaSPARHA and AcaSPARHA (left side, top and bottom) and of tetrameric complexes of HNH domains of MllaSPARHA and AcaSPARHA with substrate dsDNA (right side, top and bottom). The PAE values show the expected position error at residue x if the predicted and true structures were aligned on residue y. The positions of individual domains of the SPARHA complexes along the model coordinates are shown at the bottom of each plot. The color gradient from blue to red indicates the error value, from low to high. The alignment of the HNH domain relative to the rest of the complex is less confident in MllaSPARHA indicating that it may more flexible even in the absence of guides and targets. All graphs were created by AlphaFold Server ^63^.

## Methods

### Bioinformatic analysis of short pAgos and HNH nucleases

For the construction of the phylogenetic tree of APAZ-containing proteins, a search was performed using DELTA-BLAST (part of the BLAST 2.15.0+ package) against the nr (non-redundant proteins) RefSeq database with the following parameters: -taxids 2,2157 (Bacteria, Archaea), -num_iterations 3, -evalue 0.005.The input file contained the APAZ domains of the following proteins: WP_141644551.1, WP_215874058.1, WP_075600999.1, WP_033317603.1, WP_011516871.1, WP_046756489.1, WP_010942011.1, WP_235024190.1, WP_006053074.1, WP_109649956.1, WP_092459739.1. The identified proteins were filtered using a custom Python script, resulting in 1546 unique proteins with WP identifiers. Their sequences, as well as the sequences of neighboring proteins in the genomes, were obtained using another custom Python script via Entrez Efetch and were then annotated using rpsblast (part of the BLAST 2.15.0+ package) and InterProScan 5.72-103.0. Protein sequences for which neighboring proteins containing the PIWI domain were extracted for further analysis and clustered using MMseq2 v.16.747c6 with a sequence identity threshold of 45%. After clustering, the previously studied proteins and the proteins from this study (identifiers listed above) were added to the resulting sequences, and a total of 321 sequences were aligned using MAFFT v.7.526 (method E-INS-i). The alignments were then trimmed with trimAl v.1.5 using the parameters -gt 0.5 (gap threshold), -resoverlap 0.65, - seqoverlap 65, resulting in 272 sequences. The phylogenetic tree was constructed using IQ-TREE v.2.3.6 with the ModelFinder algorithm (which identified Q.pfam+F+R7 as the best-fit model) and ultrafast bootstrap with 1000 replicates, followed by visualization using iTOL.

For the construction of the phylogenetic tree of HNH domain-containing proteins, a search was performed using DELTA-BLAST (part of the BLAST 2.15.0+ package) against the nr (non-redundant proteins) RefSeq database with the following parameters: -taxids 2,2157 (Bacteria, Archaea), -num_iterations 5, -evalue 0.005. The input file contained the sequences of HNH domains from various defence systems (Septu, Retron, Cas9, IscB, Zorya II, CRISPR I-F, EVE, SRA, Cap5 CBASS, Vvn, homing endonuclease). The identified proteins were filtered as described above, resulting in 8412 unique proteins with WP identifiers. Their sequences were obtained and clustered as described above. After clustering, the previously studied HNH domain-containing proteins (identifiers listed above) were added to the resulting sequences, and a total of 727 sequences were aligned as described above and trimmed with trimAl v.1.5 (-gt 0.5), resulting in 724 sequences. The phylogenetic tree was constructed and visualized as described above (the selected best-fit model was VT+I+R9). Protein annotation was performed using rpsblast, InterProScan 5.72-103.0, HHsuite v. 3.3.0 with the Pfam-A database, AlphaFold 3 and the DALI server, as well as analysis of their genomic neighbors.

### Cloning, expression and purification of SPARHA

The coding sequences for AcaAgo, MllaAgo, LheAgo, and their partner proteins HNH-APAZ were codon-optimized for *E. coli* expression, obtained by custom gene synthesis and cloned into the pBAD-HisB expression plasmid downstream of the araBAD promoter with a ribosome binding site between each pair (pAgo and HNH-APAZ), resulting in the pBAD_AcaSPARHA, pBAD_MllaSPARHA, and pBAD_LheSPARHA plasmids. Site-directed mutagenesis was performed to obtain substitutions in the HNH domain (H53A for MllaSPARHA and N71A/H80A for AcaSPARHA). For protein expression, *E. coli* BL21(DE3) cells harboring pBAD_SPARHA plasmids were grown in LB medium with ampicillin (150 μg/ml) at 37 °C with shaking (200 rpm) to OD_600_ ∼0.5, followed by cold shock at 0 °C for 45 min. Protein expression was induced with 0.05% L-arabinose and carried out overnight at 16 °C. Then, the cells were harvested by centrifugation and used for protein purification.

To purify the SPARHA complexes, cells were lysed by sonication in buffer containing 20 mM HEPES, pH 8.0, 1 M KCl, 10 mM K_2_HPO_4_, 5% glycerol, 2 mM DTT, 2 mM PMSF and then centrifuged at 25,500 g for 25 min at 4 °C. The supernatant was transferred onto a HisTrap HP 1 ml column (Cytiva) pre-equilibrated with buffer containing 20 mM HEPES, pH 8.0, 0.5 M KCl, 5% glycerol, washed with the same buffer and then with the buffer containing 21 mM imidazole, and the proteins were eluted with the buffer containing 300 mM imidazole. The eluate was supplemented with EDTA (up to 5 mM) and diluted 20-fold with buffer containing 20 mM HEPES, pH 8.0, 5% glycerol, 2 mM EDTA and loaded onto a HiTrap Heparin HP column 5 ml (Cytiva) pre-equilibrated with buffer containing 20 mM HEPES, pH 8.0, 30 mM KCl, 5% glycerol and 2 mM EDTA. After washing, the complexes were eluted with a KCl gradient 30 to 800 mM in buffer containing 20 mM HEPES, pH 8.0, 5% glycerol, and 2 mM EDTA. For AcaSPARHA and LheSPARHA, the elution profiles revealed peaks at 24 mS, and for MllaSPARHA, a peak at 40 mS. The corresponding peak fractions were collected, combined, and diluted with buffer containing 20 mM HEPES, pH 8.0, 5% glycerol, 2 mM EDTA to reduce the KCl concentration to 36 mM (28 mM for LheSPARHA). Subsequently, the samples were loaded onto a MonoQ 1 ml column (Cytiva) pre-equilibrated with buffer containing 20 mM HEPES, pH 8.0, 35 mM KCl (25 mM for LheSPARHA), 5% glycerol and 2 mM EDTA. After washing, the complexes were eluted with a KCl gradient from 35 mM (25 mM for LheSPARHA) to 800 mM in buffer containing 20 mM HEPES, pH 8.0, 5% glycerol and 2 mM EDTA. The elution profile revealed two peaks for AcaSPARHA (20 and 40 mS) and LheSPARHA (8 and 45 mS). For MllaSPARHA, a single peak at 22 mS was observed. Analysis of fractions by SDS-PAGE and mass-spectrometry showed that the peaks at 20 mS and 22 mS corresponded to complexes of AcaSPARHA and MllaSPARHA, respectively. In the case of LheSPARTA, the peak at 8 mS corresponded to LheSPARHA, and the peak at 45 mS contained only LheHNH-APAZ. The fractions containing AcaSPARHA, MllaSPARHA, LheSPARHA, and LheHNH-APAZ were concentrated with an Amicon 30 kDa. The proteins were stored in a buffer with 50% glycerol and 2 mM DTT for further experiments.

### Analysis of *in vitro* activities of SPARHA

For the analysis of nuclease activity of SPARHA on linear collateral substrates, 500 nM SPARHA (MllaSPARHA, AcaSPARHA, or their CD mutants, as well as LheSPARHA or LheHNH-APAZ) were incubated with 200 nM gRNA (unless otherwise indicated) in a buffer containing 20 mM HEPES, pH 8.0, 5 μg/ml BSA, 0.4 mM DTT, 5 mM Mn^2+^ (unless otherwise indicated), 5 mM KCl for AcaSPARHA and 100 mM KCl for MllaSPARHA or LheSPARHA (unless otherwise indicated) for 15 minutes at 30 °C (unless otherwise indicated). Then, 200 nM complementary tDNA (unless otherwise indicated) was added, and after incubation for 10 minutes at 30 °C, 100 nM collateral substrate (ssDNA, RNA, dsDNA with pre-annealed strands) was added. After 1 hour (or another time as indicated in the Figures), the reaction was stopped by adding an equal volume of loading buffer containing 8 M urea, 20 mM EDTA, and 2x TBE. The reaction products were separated by 19% urea PAGE and visualized by SYBR Gold staining or fluorescence scanning (FAM for RNA substrates, HEX for dsDNA substrates) using a Typhoon FLA 9500 scanner (GE Healthcare). In the case of plasmid or genomic DNA as a substrate, the same protocol was used, except that the concentration of plasmid DNA and genomic DNA was 2 nM and 100 ng, respectively, and the reaction was stopped by adding 1/10 volume of loading buffer containing 50% glycerol, bromophenol blue, SYBR Gold (0.2 μl per 200 μl), and 0.1% SDS. The reaction products were separated on a 1% agarose gel and visualized as described above.

To analyze the multi-turnover cleavage and substrate preferences of the SPARHA complex, 500 nM MllaSPARHA was incubated at 30 °C with 500 nM gDNA and tDNA as described above. Subsequently, 5 μM of various collateral substrates (DNA, RNA, or dsDNA of identical sequences) were added. The reaction was stopped as described above at specific time points (indicated in Figure ED2b). Reaction products were separated by 19% urea PAGE and visualized by fluorescence scanning (FAM for the RNA substrate) or phosphorimaging for ssDNA and dsDNA (labeled with ^32^P at the 5’-end) using a Typhoon FLA 9500 scanner (GE Healthcare). See Table S2 for oligonucleotide sequences.

### Fluorescence assay

The AcaSPARHA complex (1 μM) was incubated with ori-gRNA-20nt (1.3 μM) for 15 minutes at 30 °C, then cooled on ice for 5 minutes. The pre-annealed collateral substrate dsDNA-32nt (1 μM) labeled with the ROX fluorophore and BHQ2 quencher was added on ice. Subsequently, ori-tDNA-20nt was added at various concentrations (as indicated in the legend to ED4b) on ice. Aliquots (15 μL) were transferred into a pre-chilled 384-well plate (CORNING, low volume), which was placed into a CLARIOstar reader. The incubation was carried out at 30 °C, and fluorescence measurements were taken every 5 minutes using EDR (Enhanced Dynamic Range).

### Analysis of the effects of SPARHA on plasmids and phages

To analysis of the effects of SPARHA expression on cell growth depending on the presence of interfering plasmids, *E. coli* BL21(DE3) cells were transformed with empty pBAD or pBAD_SPARHA plasmids (AcaSPARHA; LheSPARHA; or MllaSPARHA including its mutants; all Amp^R^) and a second plasmid, either pCDFDuet-1 with the GST gene insertion (Sm^R^) or pACYC184 (Cm^R^). For analysis of growth curves, overnight cultures (grown with the corresponding antibiotics and 0.5% glucose) were diluted 75-fold in LB containing Amp, and 200 μL of the diluted culture was mixed with 40 μL of L-Ara to a final concentration of 0.2%. Subsequently, the cells were incubated at 25 °C (or 31.5 °C) with shaking in a 48-well plate using a CLARIOstar reader, with OD_600_ measured every 10 minutes. To determine the number of viable cells, samples were collected from the plate (after 8 hours of incubation at 25°C or after 10.5 hours of incubation at 31.5°C), and 10-fold serial dilutions were prepared and plated on LB agar containing Amp or the antibiotic corresponding to the interfering plasmid, with the addition of 1% glucose to suppress SPARHA expression. The number of colonies was counted after overnight incubation of the plates at 37°C. For the experiment shown in ED5f, the protocol described above was used, except that antibiotics were not added to the overnight cultures or during the dilution of overnight cultures and after collecting samples from the 48-well plate, their 10-fold serial dilutions were plated on LB agar plates either without antibiotics or with the antibiotics corresponding to the plasmids.

For cell staining, *E. coli* BL21(DE3) containing empty pBAD, MllaSPARHA, and CD mutant of MllaSPARHA in the presence of the second plasmid pCDF were collected from a 48-well plate after 7.5 hours of incubation at 25 °C with shaking in a CLARIOstar reader. The cells were pelleted by centrifugation, washed with 0.9% NaCl, and then resuspended in 0.9% NaCl to OD_600_ ∼2.5. An equal volume of a mixture of SYTO9 (excitation/emission maxima: 485/500 nm) and SynaptoProbe Red (analogue of FM4-64; excitation/emission maxima 515/640 nm) in 0.9% NaCl was added to the cells to final concentrations of 1 μM each. After 10 minutes of incubation at room temperature, 4 μL of the stained cells were visualized using a ZEISS LSM 900 confocal laser scanning microscope (Carl Zeiss) with excitation wavelength of 488 nm for SYTO9 and for SynaptoProbe Red, and images were taken using a 63x oil immersion objective. The experiment was performed in three biological replicates, with 5-10 fields per replicate.

To identify the effects of SPARHA on phage infection, the efficiency of plating (EOP) was determined*. E. coli* BL21(DE3) cells were transformed with either the empty pBAD or MllaSPARHA (WT or CD). Overnight cultures were diluted 30-fold (200 μL into 6 mL) in LB containing Amp and 0.2% L-Ara, and then incubated at 30 °C with shaking until OD_600_ ∼0.5. Then, 0.5 mL of cells were mixed with 32 mL of 0.4% top agar containing 10 mM MgCl_2_, 5 mM CaCl_2_, Amp and 0.1% L-Ara. PFU determination was performed on plates containing bottom 1.5% agar and top agar (as described above), onto which 10 μL of 10-fold serial dilutions of phages HK140, HK243, mEp243, HK578, HK75, P1, λ, T7, T5, T6, HK225, HK106 and HK446 were spotted. The plates were incubated overnight at 25 °C or 30 °C (in the case of incubation at 30 °C, the plates contained 0.2% L-Ara). The PFU numbers were counted after 18 hours for phages HK140, HK243, and mEp243, and after 20 hours for phage HK578. Additionally, plates with phages HK140 and HK243 were photographed after 14 hours of incubation at 25 °C (as shown in ED7d). In all cases, the incubation time is indicated above the plates.

### Analysis of small RNAs associated with SPARHA

To isolate small RNAs associated with MllaSPARHA in the presence of the interfering plasmid pCDF, overnight culture of *E. coli* BL21(DE3) cells transformed with pBAD_MllaSPARHA and pCDF-duet plasmids was added to LB medium containing Amp (150 μg/mL) and incubated at 30 °C with shaking for 1.5 hours. Subsequently, L-Ara was added to a final concentration of 0.2%, and the culture was grown at 30 °C with shaking for 6.5 hours. The cells were harvested by centrifugation (20 min, 5000 rpm, 4 °C), resuspended in lysis buffer (20 mM HEPES, pH 8.0, 1 M KCl, 10 mM K_2_HPO_4_, 2 mM PMSF, 2 mM DTT) and lysed by sonication. The supernatant was loaded onto a HisTrap HP 1 ml column (Cytiva) pre-equilibrated with buffer (20 mM HEPES, pH 8.0, 1 M KCl, 5% glycerol). After washing, the protein complex was eluted with the same buffer containing 300 mM imidazole. The eluate was concentrated using an Amicon 30 kDa device and loaded into 15% urea PAGE. The gel was visualized using SYBR Gold staining with a UV transilluminator, and the small RNAs were extracted from the gel, followed by overnight elution in 0.4 M NaCl.

For isolation of small RNAs associated with AcaSPARHA in the presence of the interfering plasmid pCDF, overnight culture of cells transformed with pBAD_AcaSPARHA and pCDF-duet plasmids was added to LB medium containing Amp (150 μg/mL) and 0.1% L-Ara, then grown for 8 hours at 30 °C with shaking. The cells were harvested by centrifugation, resuspended in lysis buffer (20 mM HEPES, pH 8.0, 0.5 M KCl, 10 mM K_2_HPO_4_, 2 mM PMSF, 2 mM DTT) and lysed by sonication. Protein isolation of AcaSPARHA was performed as described above including heparin and MonoQ chromatography steps, with the exception that EDTA was not added to the eluate after Ni-chelating chromatography and was excluded from all buffers. Peak of AcaSPARHA associated with small RNAs was eluted at 37 mS from the MonoQ column. The collected fractions were used for RNA isolation via phenol-chloroform extraction.

In both cases, small RNAs were precipitated using 3 M ammonium acetate, 96% ethanol, and glycogen, ethanol-washed, dissolved in water, phosphorylated using ATP and T4 polynucleotide kinase (NEB), and again precipitated, resulting in the final purified small RNAs. RNA libraries were prepared using the NEBNext Small RNA Library Prep Set for Illumina (Multiplex Compatible). Both RNA libraries (two biological replicates for MllaSPARHA and AcaSPARHA) were sequenced using Illumina HiSeq2500 (single-end 50-nucleotide reads). The list of sequenced small RNA libraries with corresponding accession numbers is shown in Table S3.

Raw reads were processed to remove adapters using Trim Galore v.0.6.10, and the sequences were trimmed to the length of 15 to 25 nucleotides with Seqtk v.1.4-r122. Quality control before and after trimming was performed using FastQC v.0.12.1. The resulting libraries were mapped to the *E. coli* BL21(DE3) genome (NC_012971.2) and plasmids (pBAD_MllaSPARHA or pBAD_AcaSPARHA and pCDF) using STAR v.2.7.11b (with 97% alignment identity and intron exclusion, MatchNmin 14). Primary reads were extracted and used to generate sequence logos (16–25 nt or ≥20 nt in length) with WebLogo. Read counting of primary reads mapped to genes was performed using featureCounts v.2.0.8 (-s 1 for sense strand, -s 2 for antisense strand). The resulting values were normalized using RPKM, and distribution plots were created using custom Python scripts.

### Mapping of the sites of DNA cleavage by SPARHA

To analyze the cleavage specificity, MllaSPARHA (500 nM) was preincubated with UA-gRNA-18nt (200 nM) in a reaction buffer (20 mM HEPES pH 8.0, 100 mM KCl, 5 μg/ml BSA, 2 mM DTT, 5 mM MnCl_2_) for 15 minutes at 30 °C. Control sample without MllaSPARHA was obtained in parallel. Complementary target DNA (200 nM) was added to the reaction mix and the samples were incubated for 10 minutes at 30 °C. Genomic DNA (80 ng) was added (purified from *E. coli* BL21(DE3) transformed with pBAD_Mlla and pCDF-Duet) and the cleavage reactions were performed for 10 minutes at 30 °C. The samples were mixed with 10^x^ loading buffer (50% glycerol, 0.1% SDS, SYBR Gold), chilled in ice, and aliquots were analyzed by 1 % agarose gel electrophoresis to confirm DNA smearing in the reaction containing MllaSPARHA. DNA from the remaining samples was treated with phenol-chloroform and ethanol-precipitated. For library preparation, DNA ends were blunted using the DNA end repair mixture and adapter sequences described previously ^64^. The samples were treated with 2 μl of 3 U/μl T4 DNA polymerase (NEB), 2.5 μl of 10 U/μl T4 polynucleotide kinase (NEB), 0.5 μl of 5 U/μl, Large Klenow Fragment of DNA Polymerase I (NEB) in total volume of 50 μl containing 5 μl 10^x^ T4 DNA ligase buffer (NEB) and 2.5 μl 10 mM dNTP mix (Thermo Fisher Scientific, R0191) for 45 min at 25 °C. DNA was purified with Agencourt AMPure XP beads and eluted with 25 μl of 10 mM Tris-HCl (pH 8.0). To perform the A-tailing reaction, the DNA samples were supplemented with 15 μl of MiliQ-grade water, 5 μl of 10^x^ NEBuffer 2.1 (B6002S), 2.5 μl of 10 mM dATP and 2.5 μl of 5 U/μl Klenow (exo-) (NEB). The reactions were carried out for 45 min at 37 °C and the enzyme was heat-inactivated by incubation for 20 min at 65 °C. DNA was purified using Agencourt AMPure XP beads and eluted with 40 μl of 10 mM Tris-HCl (pH8.0).

Detector adapter with a 3’-dT overhang was generated by annealing Detector_adapter_dir and phosphorylated Detector_adapter_rev oligonucleotides as described previously ^64^. The resulting adapter (2.5 μl of 10 μM solution) was ligated to the purified 3’-dA overhanged cleavage products in 50 μl samples containing 5 μl of 1 U/μl T4 DNA ligase and 5 μl of 10^x^ T4 DNA ligase buffer (NEB) for 16 h at 22 °C. The ligated DNA was purified using Agencourt AMPure XP beads and eluted with 28 μl of 10 mM Tris-HCl (pH 8.0). Next, sequencing primers were added using the NEBNext Ultra II FS DNA Library Prep Kit for Illumina. The resulting samples were amplified with dual index primers (NEBNext Multiplex Oligos for Illumina) and purified with Agencourt AMPure XP beads. PCR products were sequenced on a NovaSeq 6000 System (Illumina) in the 100 bp paired-end mode. The list of sequenced DSB libraries with corresponding accession numbers is shown in Table S3.

Raw reads were processed to filter by quality using Trim Galore v.0.6.10; quality control before and after filtering was performed with FastQC v.0.12.1. Subsequently, reads containing the detector adapter sequence at the 5’-end were extracted from libraries for further analysis (for MllaSPARHA and control) and the adapter sequence was further removed, using custom Python scripts. The resulting library was mapped to the genome (NC_012971.2) and plasmids (pBAD_MllaSPARHA and pCDF-Duet) using STAR v.2.7.11b (with 97% alignment identity and intron exclusion).

To construct sequence logos, the first 5 nucleotides of primary reads (distance from +1 to +5, Fig. 4c, ED8b) and 6 nucleotides to the left of the read start (distance from -5 to 0, Fig. 4c, ED8b) were extracted from the reference genomic and plasmid sequences using a custom Python script. Sequence logos of the resulting FASTA files were generated using the WebLogo. To analyze the distribution statistics of distances between the 5’-ends of reads on the + and – strands in the genome, a custom Python script based on a binary search algorithm was used. The resulting statistics were visualized with a custom Python script. To construct a sequence logo for pairs with a distance of 1, reads from the identified pairs were extracted into a separate BAM file, and the logo was generated as described above.

### Size-exclusion chromatography and SEC-MALS

The absolute molar masses and the oligomeric state of the apoforms of AcaSPARHA and MllaSPARHA were determined by size-exclusion chromatography (SEC) coupled to multi-angle light scattering (MALS). Proteins were pre-incubated for at least 15 min at 10 °C in the Vialsampler (G7129A, Agilent) and loaded on a Superdex 200 Increase 10/300 column (Cytiva) operated using an Agilent 1260 Infinity II chromatography system equipped with a 1260 Infinity II WR diode-array detector (G7115A, Agilent), a miniDAWN detector (Wyatt Technology) and a refractometric detector (G7162A, Agilent, operated at 35 °C). The column buffer contained 20 mM HEPES-KOH pH 8.0 with 100 mM KCl, and the flow rate was 1 ml/min. Post-column filters were not used. The elution profiles were followed by absorbance at 280 nm and changes of the refractive index, as well as static light scattering at three angles. The data were collected and processed in Astra 8.0 software (Wyatt Technology) using an refractometric detector as a concentration source (dn/dc was taken equal to 0.185 for each protein) and 250,000 attenuation of the output RI signal. Normalization of the static light scattering signals at different angles detected by miniDAWN was done using a pre-run of a BSA standard (Wyatt). The data were analyzed in ASTRA 8.0 (Wyatt Technology, USA) using dn/dc = 0.185. The polydispersity index (the *M*_w_/*M*_n_ ratio) of the protein peaks was around 1.000.

To screen for the effects of the metal ion, RNA guide or DNA target on the apparent size and oligomeric behavior of protein samples, the samples were pre-incubated at 30 °C within the Vialsampler and injected in 28 μl aliquots onto a Superdex 200 Increase 5/150 column (Cytiva) operated at a 0.45 ml/min flow rate. The column was pre-equilibrated with 20 mM HEPES-KOH pH 7.0, containing 50 mM KCl and 0.1 mM MnCl_2_. In the presence of guide RNA and target DNA, both wild-type and CD AcaSPARHA displayed a large peak in the void column volume, which for Superdex 200 Increase corresponded to ≥600 kDa. To assess more specifically what large size this species corresponds to, we performed SEC-MALS using a Superose 6 Increase 10/300 column (Cytiva), which has an exclusion limit of 5 MDa. Column buffer was 20 mM HEPES-KOH pH 7.0 containing 50 mM KCl and 0.1 mM MnCl_2_.

The flow rate was 1 ml/min. The data were analyzed in ASTRA 8.0 (Wyatt Technology, USA) using dn/dc = 0.185 and the extinction coefficient ε (0.1%) at 280 nm of 1.29 (mg/ml)^-1^cm^-1^. The protein conjugate analysis implemented in Astra 8.0 (Wyatt) was used to assess mass contributions from protein and bound nucleic acid components in the elution peaks, using the dn/dc and ε (0.1%) values indicated above for Aca, dn/dc = 0.17, ε (0.1%) at 280 nm of 15.8 (mg/ml)^-1^cm^-1^ for the guide RNA (AUUUGGUUGCGGUGCUCUGC), or dn/dc = 0.17, ε (0.1%) at 280 nm of 10 (mg/ml)^-1^cm^-1^ for the RNA/DNA duplex (AUUUGGUUGCGGUGCUCUGC/GCAGAGCACCGCAACCAAAT).

### Transmission electron microscopy with negative staining

To visualize the SPARHA complex with gRNA and tDNA by negative-stain transmission electron microscopy, 2.5 μM complexes of MllaSPARHA, AcaSPARHA, or their CD mutants were incubated with 2.5 μM gRNA in a buffer containing 20 mM HEPES (pH 7.0), 100 μM Mn^2+^, and either 100 mM KCl for WT and CD MllaSPARHA or 50 mM KCl for WT and CD AcaSPARHA for 10 min at 30 °C. Subsequently, 2.5 μM complementary tDNA was added, and incubation was continued for additional 2.5 min. To prepare the samples of SPARHA for TEM, 3 µl of sample suspension was applied to a carbon substrate film (EMCN, China) hydrophilized with glow discharge. Negative staining was performed using 1% uranyl acetate for 30 seconds.

TEM imaging was carried out using JEM-1400 120 kV transmission electron microscope (JEOL, Japan) equipped with Rio 9 camera (Gatan, USA). For 3D reconstruction, image series were acquired with JEM-2100 200 kV transmission electron microscope (JEOL, Japan) equipped with a DE-20 detector (Direct Electron, USA). Automated data collection was performed with SerialEM ^65^ at a defocus of -1.8 µm. For each sample, 600-1200 images were acquired. The contrast transfer function (CTF) was estimated with CTFFIND4 ^66^. 3D reconstruction was performed in cryoSPARC ^67^. Individual helical filament segments (particles) were picked using template-based selection based on manually selected particles.

Structural heterogeneity, particularly the presence of different helix types within the Mlla protein complex, was resolved through multiple rounds of 2D classification. Helix parameters were initially determined via asymmetric helical refinement and subsequently refined in the final round of helical refinement. The resulting reconstructions were generated from 3,000– 12,000 particles, depending on the sample, and achieved a resolution of 16 Å based on the FSC 0.143 criterion. This resolution is consistent with the limitations of the negative staining method.

### Structural modeling

All models were predicted by AlphaFold 3 (AF3). The model of the Mlla HNH tetramer with dsDNA was manually adjusted using PyMol, based on the interactions of residues between HNH dimers in the structure of the Cap5 tetramer (PDB ID:8FMF). For building the models of oligomers of AcaSPARHA and MllaSPARHA containing gRNA and tDNA, the tetramer models were copied and aligned head-to-tail by the MllaSPARHA or AcaSPARHA chains using UCSF ChimeraX.

### Statistical analysis

Statistical analysis of the experimental data was performed with the statannotations Python package, with P-values calculated via an independent samples t-test and corrected using either the Bonferroni or Holm-Bonferroni correction, as indicated in the figure legends for each case.

## Acknowledgements

We thank Maria Prostova for the search and selection of SPARHA systems, Maria Prostova, Daria Gelfenbeyn, Vladimir Panteleev, Alena Zavgorodniaia and Alexei Aravin for invaluable advices and discussions. The transmission electron microscopy studies were carried out at the Shared Research Facility “Electron microscopy in life sciences” at Moscow State University (Unique Equipment “Three-dimensional electron microscopy and spectroscopy”). This work was supported by the Russian Science Foundation (grant 22-14-00182 to A.Kulbachinskiy).

## Data and Code Availability

The data generated during this study are included in the published article. The guide RNA sequencing and DNA cleavage datasets generated in this study are available from the Sequence Read Archive (SRA) database. The code used for data analysis is available at the GitHub repository. All primary data are available from the corresponding authors upon request.

## Author contributions

Conceptualization, supervision, experimental design: A.Kanevskaya and A.Kulbachinskiy. Phylogenetic analysis, structural modeling, molecular cloning, protein purification, experiments *in vitro*, plasmid interference, phage infection, fluorescence microscopy, preparation and analysis of small RNA libraries, analysis of DSB DNA libraries: A.Kanevskaya. Mapping of DSB cleavage sites by SPARHA: L.Lisitskaya and A.Kanevskaya. SEC-MALS analysis: N.N.Sluchanko and A.Kanevskaya. Electron microscopy: A.V.Moiseenko, A.Kanevskaya, O.S.Sokolova. Preparation of Figures: L.Lisitskaya and A.Kanevskaya with contribution from all of the authors. Writing - original draft: A.Kulbachinskiy. Writing - review and editing: A.Kulbachinskiy and A.Kanevskaya with contribution from all of the authors.

## Competing interests

The authors declare no competing interests.

**Correspondence and requests for materials** should be addressed to A.Kanevskaya or A.Kulbachinskiy.

